# The druggable genome and support for target identification and validation in drug development

**DOI:** 10.1101/066027

**Authors:** Chris Finan, Anna Gaulton, Felix. A Kruger, Tom Lumbers, Tina Shah, Jorgen Engmann, Luana Galver, Ryan Kelley, Anneli Karlsson, Rita Santos, John P. Overington, Aroon D. Hingorani, Juan P. Casas

## Abstract

Target identification (identifying the correct drug targets for each disease) and target validation (demonstrating the effect of target perturbation on disease biomarkers and disease end-points) are essential steps in drug development. We showed previously that biomarker and disease endpoint associations of single nucleotide polymorphisms (SNPs) in a gene encoding a drug target accurately depict the effect of modifying the same target with a pharmacological agent; others have shown that genomic support for a target is associated with a higher rate of drug development success. To delineate drug development (including repurposing) opportunities arising from this paradigm, we connected complex disease- and biomarker-associated loci from genome wide association studies (GWAS) to an updated set of genes encoding druggable human proteins, to compounds with bioactivity against these targets and, where these were licensed drugs, to clinical indications. We used this set of genes to inform the design of a new genotyping array, to enable druggable genome-wide association studies for drug target selection and validation in human disease.

## Introduction

Only 4% of drug development programmes yield licensed drugs *(1, 2)*, largely because of two unresolved systemic flaws: (1) preclinical experiments in cells, tissues and animal models and early phase clinical testing to support drug target identification and validation are poorly predictive of eventual therapeutic efficacy; and (2) definitive evidence on the validity of a new drug target for a disease is delayed until late phase development (in phase II or III randomised controlled trials; RCTs). Reasons for poor reliability of preclinical studies include suboptimal experimental design with infrequent use of randomisation and blinding *(3)*; species differences; inaccuracy of animal models of human disease *(4, 5)*; and over-interpretation of nominally significant experimental results *(6–8)*. Human observational studies can mislead for reasons of confounding and reverse causation. Evidence on target validity from phase I clinical studies can also be inadequate (since phase I studies primarily investigate pharmacokinetics and tolerability, are typically small in size, of short duration and measure a narrow range of surrogate outcomes, often of uncertain relevance to perturbation of the target of interest) *(9)*. Since the target hypothesis advanced by preclinical and early phase clinical studies is all too frequently false, expensive late-stage failure in RCTs from lack of efficacy is a common problem affecting many therapeutic areas *(10)*, posing a threat to the economic sustainability of the current model of drug development.

Genetic studies in human populations imitate the design of an RCT without requiring a drug intervention *(11–13)*. This is because genotype is determined by a random allocation at conception according to Mendel’s second law (Mendelian randomisation - MR) *(12, 14)*. Single nucleotide polymorphisms (SNPs) acting in *cis* (i.e. variants in or near a gene that associate with the activity or level of the encoded protein) can therefore be used as a tool to deduce the effect of pharmacological action on the same protein in an RCT. Numerous proof of concept examples have now been reported *(15, 16, 11, 17, 13, 18, 19)*, including the striking correlation between the association of 80 circulating metabolites with a SNP in the *HMGCR* gene that encodes the target for statin drugs, and the effect of statin treatment on the same set of metabolites *(20)*. SNPs acting in *cis* are a general feature of the human genome *(21)*; and population and patient datasets with stored DNA and genotypes linked to biological phenotypes and disease outcome measures are now widely available for this type of study.

By extension, disease-associated SNPs identified by GWAS could be re-interpreted as an under-utilised source of randomised human evidence to aid drug target identification and validation. For example, loci for type-2 diabetes identified by GWAS include genes encoding targets for the glitazone and sulphonylurea drug classes already used to treat diabetes *(22, 23)*. Apparently sporadic observations such as this suggest that numerous, currently unexploited disease-specific drug targets should exist among the thousands of other loci identified by GWAS and similar high quality genetic association studies. Recent studies of advanced or completed drug development programmes (mostly based on established approaches to target identification) have also indicated that those with incidental genomic support had a higher rate of developmental success *(24–27)*.

Fulfilling the potential of GWAS (and studies using disease-focused genotyping arrays) for drug development requires mapping disease- or biomarker-associated SNPs to genes encoding druggable proteins and to any allied drugs and drug-like compounds. The set of proteins with potential to be modulated by a drug-like small molecule has been predicted on the basis of sequence and structural similarity to the targets of existing drugs, the set of encoding genes being referred to as the druggable genome. Hopkins and Groom identified 130 protein families and domains found in targets of drug-like small molecules known at the time, and over 3000 potentially druggable proteins containing these domains *(28)*. A similar estimate was made by Russ and Lampel, using a later human genome build *(29)*. Kumar et al. utilized these privileged protein families (plus other families of particular relevance to cancer) to manually curate lists of druggable proteins for inclusion in the dGene data set *(30)*. More recently, the Drug-Gene Interaction database (DGIdb) has been developed *(31)*, which integrates data from each of the previous efforts together with a recently compiled list of drug candidates and targets in clinical development *(32)* as well as information from the PharmGKB *(33)*, Therapeutic Target Database (TTD) *(34)* and DrugBank *(35)* databases.

However, earlier estimates of the druggable genome predated contemporary genome builds and gene annotations, and also did not explicitly include the targets of bio-therapeutics, which formed more than a quarter of the 45 new drugs approved by the FDA’s Center for Drug Evaluation and Research in 2015 *(36)*, reflecting their increasing importance in pharmaceutical development. We therefore updated the set of genes comprising the druggable genome. We then linked GWAS findings curated by the National Human Genome Research Institute (NHGRI) and European Molecular Biology Laboratory–European Bioinformatics Institute (EMBL-EBI) GWAS catalog *(37)* to this updated gene set, and also to encoded proteins and associated drugs or drug-like compounds curated in the ChEMBL *(38)* and First Databank *(39)* databases. We used the linkage to explore the potential for genetic associations with complex diseases and traits to inform drug target identification and validation, as well as to repurpose drugs effective in one indication for another. Additionally, to better support future genetic studies for disease-specific drug target identification and validation, we assembled the marker content of a new genotyping array designed for high-density coverage of the druggable genome and compared this focussed array with genotyping arrays previously used in GWAS.

## Results

### Re-defining the druggable genome

We estimated 4,479 (22%) of the 20,300 protein coding genes annotated in Ensembl v.73, to be drugged or druggable. This adds 2,402 genes to previous estimates made by Hopkins and Groom or Russ and Lampel by inclusion of novel targets of first-in-class drugs licensed since 2005; the targets of drugs currently in late phase clinical development; information on the growing number of pre-clinical phase small molecules with protein binding measurements reported in the ChEMBL database; as well as genes encoding secreted or plasma membrane proteins that form potential targets of monoclonal antibodies and other bio-therapeutics. A set of 680 genes that was included in earlier estimates but not our data set consists mainly of olfactory receptors and phosphatases; both protein families have significant limitations for future exploitation as drug targets *(40, 41)* (see Figure 1 and Methods section). We stratified the druggable gene set into 3 tiers corresponding to position in the drug-development pipeline. Tier 1 (1,427 genes) included efficacy targets of approved small molecules and biotherapeutic drugs as well as clinical-phase drug candidates. Tier 2 comprised 682 genes encoding targets with known bioactive drug-like small molecule binding partners as well as those with significant sequence similarity to approved drug targets. Tier 3 contained 2,370 genes encoding secreted or extracellular proteins, proteins with more distant similarity to approved drug targets, and members of key druggable gene families not already included in Tiers 1 or 2 (GPCRs, nuclear hormone receptors, ion channels, kinases and phosphodiesterases). A full list of genes is provided in Supplementary File S1. An overview of the 15 most frequently occurring protein domain types for each tier can be found in Supplementary Table 1, based on the Pfam-A database of protein families (see Methods section Pfam-A domain content).

**Fig. 1.**
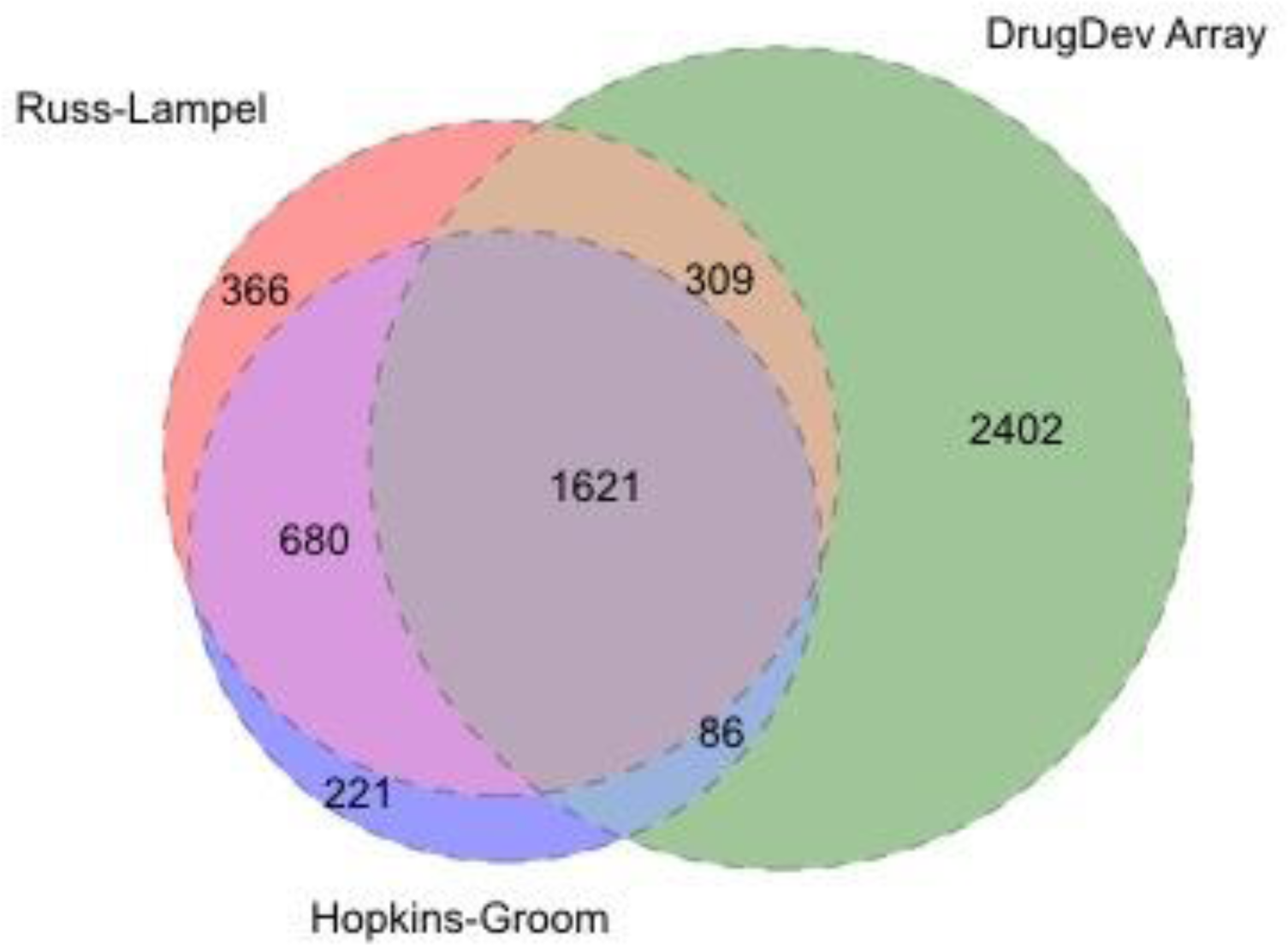
Overlap between 3 sets of druggable genes. The Venn diagram shows overlapping and distinct elements of the druggable gene sets defined by Hopkins and Groon, Russ and Lampel, and the set of druggable genes presented in this publication (DrugDev).

### Connecting loci identified by GWAS to the druggable genome

We retrieved 21,406 associations from 2,155 GWAS, of which 9,178 surpassed the significance threshold of p≤5×10^-8^ (see Methods section). The retrieved associations spanned 315 Medical Subject Heading (MeSH) disease terms, which can be stratified into twenty-four MeSH root disease areas and three MeSH Psychiatry and Psychology areas (Table 1). Variants associated with common diseases and biomarkers had median minor allele frequency 0.29 (interquartile range, IQR 0.21) based on 7,387 GWAS-significant records with risk allele frequency data, reflecting the preponderance of common variants on widely used genotyping arrays. The median odds ratio (OR) for GWAS significant studies of disease end-points was 1.24 (IQR 0.31) (based on 3,367 GWAS significant results with effect size data). We examined sequence ontology consequence types *(42)* of disease and biomarker-associated variants and found most to be non-coding, mainly intronic, presumably altering or marking variants that alter mRNA expression or availability, or marking variants that alter structure or activity of encoded proteins (Supplementary Figure S1C).

**Table 1.** Count of GWAS published per disease area.

| Mesh term | Count |
| --- | --- |
| neoplasms | 187 |
| immune system diseases | 130 |
| skin and connective tissue diseases | 107 |
| digestive system diseases | 106 |
| nervous system diseases | 104 |
| mental disorders | 85 |
| cardiovascular diseases | 84 |
| nutritional and metabolic diseases | 83 |
| endocrine diseases | 77 |
| musculoskeletal diseases | 57 |
| male urogenital disorders | 52 |
| eye diseases | 50 |
| respiratory diseases | 47 |
| haematological diseases | 43 |
| female urogenital diseases and pregnancy complications | 41 |
| pathological signs and symptoms | 34 |
| congenital disorders | 29 |
| viral diseases | 19 |
| oral diseases | 17 |
| substance-related disorders | 11 |
| diseases of the ear, nose or throat | 8 |
| parasitic diseases | 4 |
| bacterial and fungal infections | 2 |
| behavioural disorders | 1 |
| wounds and injuries | 1 |
| psychological phenomena and processes | 1 |
| occupational diseases | 1 |

**Table 2.** Number of unique GWAS associations mapping to drug targets for licensed drugs curated according to the correspondence between the GWAS association and drug indication.

| Category | # Associations | # drug targets |
| --- | --- | --- |
| Disease association and treatment indication precisely concordant* | 56 | 30 |
| Disease association and treatment indication concordant within the same disease area* | 13 | 9 |
| Disease association concordant with a biomarker of therapeutic efficacy | 97 | 37 |
| Disease association corresponding to a mechanism-based adverse effect* | 76 | 27 |
| Disease association with a known biomarker of therapeutic efficacy that can also be responsible for a mechanism-based side effect* | 32 | 8 |
| Discordant disease association and target indication considered to imply a potential repurposing opportunity | 1523 | 144 |
| Discordant disease association and target indication considered to imply either a repurposing opportunity or mechanism-based side effect depending on the direction | 108 | 52 |
| Curators unable to agree |  | 4 |

\*Refers to a target effect rediscovery (see text)

Of the 9,178 GWAS significant associations, 8,879 mapped to 5,084 unique intervals defined as containing all SNPs in linkage disequilibrium (LD) (with an r^2^ ≥ 0.5) with the SNP exhibiting the most significant association, applying an upper physical bound of 1Mb either side of this variant (see Methods section). The remaining 299 associations were either not in LD with any other variants, or not present in the 1000 genomes phase 3 panel. Such associations were assigned a nominal interval of 2.5kbp either side of the most significantly associated SNP. The frequency distribution of unique genes (and druggable genes) in LD intervals corresponding to unique significant associations were both right skewed (Figure 2), and there was a correlation between LD interval size and the number of resident genes (Supplementary Figure 3).

**Fig. 2.**
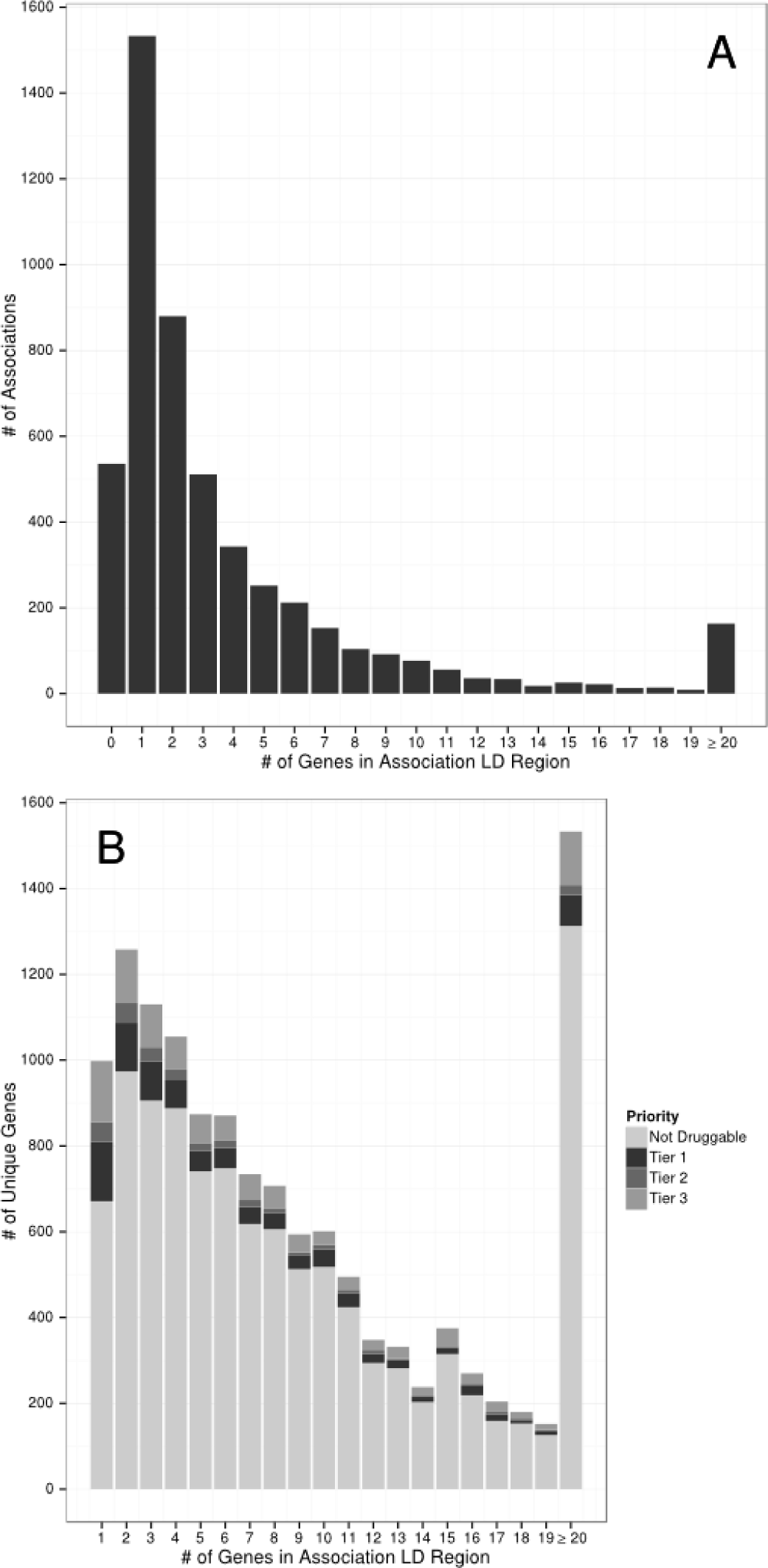
LD region summary. Part A shows the numbers of unique significant associations in the GWAS catalogue that have 0 or more genes in their LD regions. Note that there are 299 associations that had no LD region or were not present in the 1000 genomes, which are not shown in this figure. Part B, shows the number of unique genes that occupy LD regions with at least 1 gene. The counts are partitioned into genes that are not predicted to be druggable or the various druggable tiers

Of the 5,084 unique LD intervals, 1,533 (30.2%) contained a single gene. Of these, 532 also contained a single gene from the druggable set: 233 from Tier 1, 76 from Tier 2 and 223 from Tier 3. Of the remaining genomic intervals, 17.3% (880) mapped to intervals containing two genes, 10.1% (511) contained three genes 6.7% (343) contained four genes and 25.2% (1281) contained five or more genes. Additionally, 536 (10.5%) of regions had no gene in the LD interval. For the 1624 LD intervals containing two or more genes at least one of which was druggable, the median distance of the closest druggable gene to the reported GWAS variant was 4.98 kbp (IQR 37.7 kbp), where the distance was set to 0 bp for GWAS variants lying within a gene, and a druggable gene was among the two most proximal genes in 67.1 % of these LD intervals (1089) (Figure 3). We identified a total of 3052 genes in the druggable set that were not represented in any of the LD intervals corresponding to a GWAS association; 62.7%, 69.2% and 71.6% of Tier 1,2 and 3 genes respectively.

**Fig. 3.**
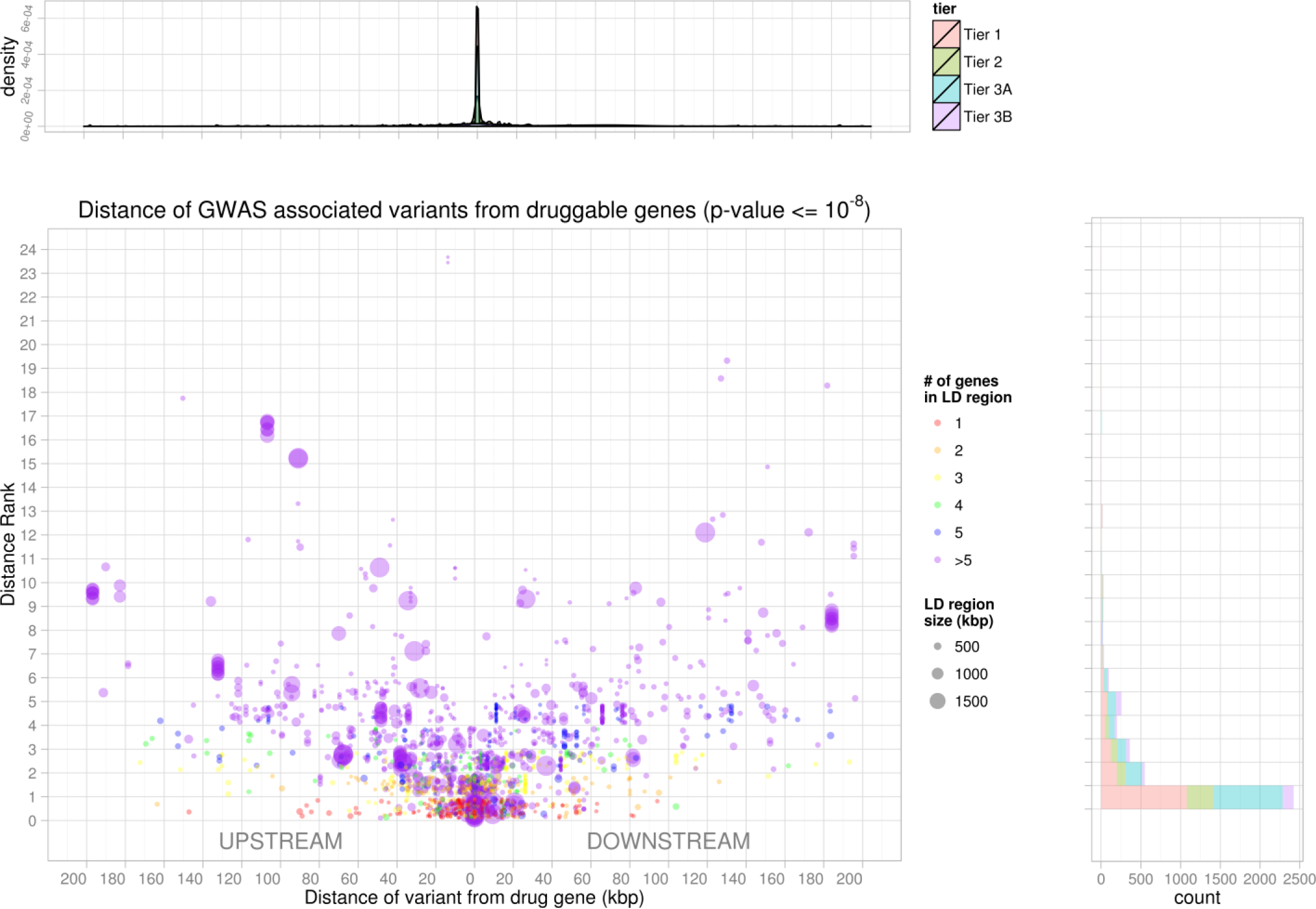
Proximity and distance rank of druggable genes to GWAS SNPs. Each point in the scatterplot corresponds to a GWAS signal located in an interval containing a druggable gene. The position on the x-axis indicates the distance of the SNP from the druggable gene. Position in the y-axis indicates the number of genes in the same interval that are closer to the signal than the druggable gene. The top panel indicates the signal density for all such SNPs, while the side panel provides the counts of signals by the distance rank of the druggable gene divided by Tier (see Methods for further details).

### Linking GWAS associations to licensed drug targets

We found that 1,291 GWAS associations defined 1,072 LD intervals containing 532 druggable genes from Tier 1, which includes the targets of licensed drugs. 479 of the intervals contained a single drug target and 593 contained two or more targets. For the set of LD intervals containing genes encoding the targets of licensed drugs, two clinically qualified curators blinded to the identity of the genes, independently evaluated the correspondence between the disease association from the GWAS and the treatment indication(s) for drug(s) acting on the target(s) encoded by a druggable gene in the interval. The curation process is described in the Methods section. Our curators identified 56 unique associations (30 unique drug targets) where the treatment indication and genetic association were precisely concordant and 13 associations (9 targets) where the indication and association came from the same disease area (e.g. a GWAS in one form of epilepsy identifying a drug target for a different form of epilepsy). 97 associations (mapping to 37 licensed drug targets) corresponded to a biomarker known to be altered by treatment with the corresponding drug (e.g. an LD interval containing the gene encoding the interleukin-6 receptor was identified in a GWAS of C-reactive protein, a biomarker known to be altered by the action of the interleukin-6 receptor blocker, tocilizumab). A further 76 associations (27 licensed drug targets) were identified through a genetic association with a mechanism-based adverse effect, e.g. in a GWAS of heart rate, the SNP rs3143709 defined an LD interval containing the gene ACHE (acetylcholinesterase) encoding the target of cholinesterase inhibitors used in the treatment of myasthenia gravis, which have the side effect of lowering heart rate *(43)*. A further 32 genetic associations (corresponding to 8 targets) were with a quantitative trait that could be either a marker of therapeutic efficacy or a mechanism-based side effect, as in the case of QT interval in the context of anti-arrhythmic drug therapy. In all, GWAS ‘rediscovered’ 74 licenced drug targets through disease indications, mechanism of action or via mechanism-based adverse effects (the numbers for the categories above are non-additive because some targets overlap categories). Illustrative examples of the curation are shown in Table 3.

**Table 3.** Illustrative examples of mapping SNPs curated in the GWAS catalogue to LD intervals containing targets of licensed and clinically used drugs. The gene encoding the drug target is listed using Human Genome Nomenclature Catalogue designation. Drug names and indications are from First Data bank. GWAS SNPs are listed according to Refseq number and physical distances are in base pairs (bp). Curation code refers to the correspondence between the treatment indication and GWAS disease or trait association (see Text). Examples are shown of treatment indication rediscoveries (Curation codes 1 and 2). For many of these the drug target gene is the sole occupant of the LD interval defined by the GWAS SNP. Examples come from a variety of disease areas and, for some diseases (e.g. type 2 diabetes and rheumatoid arthritis), multiple target rediscoveries are noted. Examples of rediscoveries of mechanism of action (curation code 3) and mechanism-based side effects are also seen (curation code 4)

| Gene | Drug | Molecule type | Curation code | GWAS EFO term | Drug Indication (FDB) | Variant | Reference (pmid) | Minimum distance from druggable gene (bp) | Distance rank of druggable gene | Genes In LD interval | Druggable genes in LD interval |
| --- | --- | --- | --- | --- | --- | --- | --- | --- | --- | --- | --- |
| **ALDH2** | DISULFIRAM | Small molecule | 1 | alcohol drinking \| drinking behavior | Alcoholism (adjunctive treatment) | rs11066280\|<br>rs12229654\|<br>rs2074356 \|<br>rs671 | 21270382\|<br>21372407\|<br>23364009\|<br>24277619 | 6016 - 790230 | 1 -18 | 22 - 33 | 2 - 4 |
| **PDE4D** | AMINOPHYLLINE | Small molecule | 1 | asthma | Acute asthma \| Acute exacerbation of chronic obstructive airways disease \| Bronchial asthma \| Chronic obstructive pulmonary disease \| Left ventricular failure - cardiac failure - cardiac asthma \| Reversible airways obstruction \| Routine maintenance therapy in chronic bronchitis and asthma | rs1588265 | 19426955 | 448153 | 1 | 2 | 1 |
| **IGF1R** | MECASERMIN | Protein | 1 | body height | Growth failure due to primary IGF-1 deficiency | rs2871865 | 20881960\|<br>25429064 | 2696 | 1 | 2 | 1 |
| **TNFSF11** | DENOSUMAB | Antibody | 1 | bone density | Prevention of skeletal related events in advanced malignancy involving bone \| Treatment of bone loss associated with hormone ablation in prostate cancer \| Treatment of osteoporosis in postmenopausal women to prevent fractures | rs17536328\|<br>rs9525638 | 24945404 | 6157 - 8295 | 1 | 1 | 1 |
| **ESR1** | TAMOXIFEN CITRATE | Small molecule | 1 | breast carcinoma | Carcinoma of breast \| Infertility - female - anovulatory | rs140068132\|<br>rs3757318 \|<br>rs9383938 | 22976474\|<br>23535729\|<br>25327703 | 9531 - 63713 | 1 - 2 | 2 | 1 |
| **PLG** | ALTEPLASE | Enzyme | 1 | coronary heart disease \| large artery stroke \| stroke | Acute ischaemic stroke: fibrinolytic treatment\| Thrombolysis in acute myocardial infarction\| Thrombolysis of occluded central venous access devices \| Thrombolytic treatment in acute massive pulmonary embolism | rs10455872 | 24262325 | 113152 | 3 | 3 | 2 |
| **TNF** | ADALIMUMAB | Antibody | 1 | Crohn's disease | Active polyarticular juvenile chronic arthritis-inadequate response to MTX \| Active progressive rheumatoid arthritis \| Moderate to severe plaque psoriasis: when other treatment is inappropriate \| Moderate/severe ulcerative colitis: when other treatment is inappropriate \| Rheumatoid arthritis when inadequate response to DMARDs incl. methotrexate \| Severe active rheumatoid arthritis \| Severe ankylosing spondylitis in adults if conventional therapy inadequate \| Treatment of active & progressive psoriatic arthritis when DMARD inadequate \| Treatment of active Crohn's disease | rs1799964 | 21102463 | 1036 | 2 | 13 | 4 |
| **CACNA1D** | AMLODIPINE | Small molecule | 1 | diastolic blood pressure | Essential hypertension when stabilised on same ingreds.in same proportions \| Hypertension-not adequately controlled by individual components \| Prinzmetal's angina \| Prophylaxis of chronic stable angina pectoris \| Treatment of essential hypertension \| | rs9810888 | 25249183 | 106912 | 1 | 1 | 1 |
| **GUCY1A3** | ISOSORBIDE DINITRATE | Small molecule | 1 | diastolic blood pressure | Angina \| Angina pectoris - prophylaxis of acute attacks \| Angina pectoris - treatment of acute attacks \| Congestive heart failure (adjunct) \| Intracoronary use during angioplasty and to prevent/relieve coronary spasm \| Left ventricular failure \| Prophylaxis of angina pectoris \| Treatment of angina pectoris \| Treatment of unresponsive LVF, either post MI or of various aetiology \| Treatment severe/unstable angina | rs13139571 | 21909115 | 7988 | 1 | 2 | 1 |
| **NPC1L1** | EZETIMIBE | Small molecule | 1 | LDL cholesterol \| low density lipoprotein cholesterol measurement \| total cholesterol measurement | Combined hyperlipidaemia: lipid lowering therapy adjunct to diet \| Homozygous familial hypercholesterolaemia (adjunct to statin therapy) \| Homozygous familial hypercholesterolaemia: Adjunct to diet \| Homozygous sitosterolaemia (phytosterolaemia) \| Primary hypercholesterolaemia (hyperlipidaemia type IIa): Adjunct to diet \| Primary hypercholesterolaemia: lipid lowering therapy adjunct to diet | rs2072183 | 20686565 \| 24097068 | 1734 | 1 | 1 | 1 |
| **PPARA** | GEMFIBROZIL | Small molecule | 1 | LDL cholesterol \| low density lipoprotein cholesterol measurement \| total cholesterol measurement | Mixed hyperlipidaemia when statin is contraindicated or not tolerated \| Primary hypercholesterolaemia: lipid lowering therapy adjunct to diet \| Reduction of cardiac events in hypercholesterolaemia \| Severe hypertriglyceridaemia with or without low HDL cholesterol | rs4253772 | 24097068 | 12050 | 1 | 7 | 2 |
| **CASR** | CINACALCET HYDROCHLORIDE | Small molecule | 1 | calcuim measurment | Homoeopathic \| Hypercalcaemia due to malignant disease \| Hypercalcaemia in primary HPT when parathyroidectomy contraindicated \| Secondary hyperparathyroidism in end stage renal disease: treatment | rs17251221 \| rs1801725 | 20661308 \| 20705733 \| 24068962 | 1585 - 12095 | 1 | 5 | 1 |
| **IL6R** | TOCILIZUMAB | Antibody | 1 | rheumatoid arthritis | Active juvenile idiopathic arthritis (unresp to NSAIDs) in comb with MTX \| Active juvenile idiopathic arthritis when inadequate response to NSAIDs \| Rheumatoid arthritis (unresp to DMARD/TNF inhib.) in comb with methotrexate \| Rheumatoid arthritis when inadequate response to DMARDs incl. methotrexate | rs2228145 | 24390342 | 14956 | 1 | 1 | 1 |
| **TNF** | ADALIMUMAB | Antibody | 1 | rheumatoid arthritis | Active polyarticular juvenile chronic arthritis-inadequate response to MTX Active progressive rheumatoid arthritis Moderate to severe plaque psoriasis: when other treatment is inappropriate Moderate/severe ulcerative colitis: when other treatment is inappropriate Rheumatoid arthritis when inadequate response to DMARDs incl. methotrexate Severe active rheumatoid arthritis Severe ankylosing spondylitis in adults if conventional therapy inadequate Treatment of active & progressive psoriatic arthritis when DMARD inadequate Treatment of active Crohn's disease | rs2596565 | 24532677 | 190015 | 24 | 145 | 27 |
| **ABCC8** | GLIMEPIRIDE | Small molecule | 1 | type II diabetes mellitus | Type 2 diabetes (NIDDM) not controlled by diet, weight loss & exercise alone | rs5219 | 19056611 | 4860 - 5802 | 3 | 5 | 3 |
| **ABCC8** | GLIPIZIDE | Small molecule | 1 | type II diabetes mellitus | Non insulin dependent diabetes mellitus when diet has failed | rs5219 | 19056611 | 4860 - 5802 | 3 | 5 | 3 |
| **ABCC8** | GLYBURIDE | Small molecule | 1 | type II diabetes mellitus | Type 2 diabetes (NIDDM) not controlled by diet, weight loss & exercise alone | rs5215 \| rs5219 | 17463248 \| 17463249 \| 19056611 \| 24509480 | 4860 - 5802 | 3 | 5 | 3 |
| **ABCC8** | NATEGLINIDE | Small molecule | 1 | type II diabetes mellitus | Control of type-2 diabetes (NIDDM) with metformin if metformin inadequate | rs5219 | 19056611 | 4860 - 5802 | 3 | 5 | 3 |
| **ABCC8** | REPAGLINIDE | Small molecule | 1 | type II diabetes mellitus | Control of type-2 diabetes (NIDDM) with metformin if metformin inadequate \| Type 2 diabetes (NIDDM) not controlled by diet, weight loss & exercise alone | rs5219 | 19056611 | 4860 - 5802 | 3 | 5 | 3 |
| **KCNJ11** | GLIMEPIRIDE | Small molecule | 1 | type II diabetes mellitus | Type 2 diabetes (NIDDM) not controlled by diet, weight loss & exercise alone | rs5219 | 19056611 | 1224 - 1306 | 1 | 5 | 3 |
| **KCNJ11** | GLIPIZIDE | Small molecule | 1 | type II diabetes mellitus | Non insulin dependent diabetes mellitus when diet has failed | rs5219 | 19056611 | 1224 - 1306 | 1 | 5 | 3 |
| **KCNJ11** | GLYBURIDE | Small molecule | 1 | type II diabetes mellitus | Type 2 diabetes (NIDDM) not controlled by diet, weight loss & exercise alone | rs5215 \| rs5219 | 17463248 \| 17463249 \| 19056611 \| 24509480 | 1224 - 1306 | 1 | 5 | 3 |
| **KCNJ11** | NATEGLINIDE | Small molecule | 1 | type II diabetes mellitus | Control of type-2 diabetes (NIDDM) with metformin if metformin inadequate | rs5219 | 19056611 | 1224 - 1306 | 1 | 5 | 3 |
| **KCNJ11** | REPAGLINIDE | Small molecule | 1 | type II diabetes mellitus | Control of type-2 diabetes (NIDDM) with metformin if metformin inadequate \| Type 2 diabetes (NIDDM) not controlled by diet, weight loss & exercise alone | rs5219 | 19056611 | 1224 - 1306 | 1 | 5 | 3 |
| **PPARG** | PIOGLITAZONE HYDROCHLORIDE | Small molecule | 1 | type II diabetes mellitus | Combination treatment of Type 2 diabetes with insulin \| Control of type-2 diabetes if metformin+sulphonylurea therapy is inadequate \| Monotherapy for type2 diabetes if overweight and metformin inappropriate \| Oral combination treatment of type 2 diabetes | rs1801282 | 24509480 | 64258 | 1 | 1 | 1 |
| **SCN1A** | OXCARBAZEPINE | Small molecule | 1 | Mesial temporal lobe epilepsy with hippocampal sclerosis \| febrile seizures | Epilepsy - combination of both partial and tonic-clonic seizures \| Epilepsy - partial seizures | rs7587026 | 24014518 | 5773 - 52194 | 1 | 3 | 1 |
| **GRIN3B** | MEMANTINE<br>HYDROCHLORIDE | Small molecule | 1 | Alzheimers disease | Moderate to severe Alzheimer's disease \| No information available | rs115550680 | 23571587 | 40689 | 8 | 8 | 2 |
| **SLC22A12** | SULFINPYRAZONE | Small molecule | 1 | urate measurement | Gout (prophylaxis) \| Gouty arthritis \| Hyperuricaemia | rs2078267 \|<br>rs478607 | 20884846 \|<br>23263486 | 23999 - 108243 | 2 - 3 | 2 - 3 | 2 |
| **SLC22A11** | PROBENECID | Small molecule | 1 | urate measurement<br>\| uric acid<br>measurement |  | rs17300741 \|<br>rs2078267 | 19503597 \|<br>20884846 \|<br>23263486 | 6233 - 8364 | 1 | 1 - 2 | 1 - 2 |
| **PDE4D** | ROFLUMILAST | Small molecule | 2 | asthma | Chronic obstructive pulmonary disease | rs1588265 | 19426955 | 448153 | 1 | 2 | 1 |
| **SCN2A** | CARBAMAZEPINE | Small molecule | 2 | febrile seizures | Epilepsy - grand mal \| Epilepsy - partial seizures \| Epilepsy - tonic-clonic seizures \| Prophylaxis of manic-depressive illness unresponsive to lithium \| Trigeminal neuralgia | rs3769955 | 25344690 | 14186 | 1 | 1 | 1 |
| **FSHR** | MENOTROPINS | Unknown | 2 | polycystic ovary syndrome | Anovulation unresponsive to clomifene citrate \| Ovarian stimulation before in vitro fertilisation \| Stimulation of spermatogenesis with concomitant hCG therapy in hypogonadism | rs2268361 | 22885925 | 12316 | 1 | 1 | 1 |
| **PLG** | ALTEPLASE | Enzyme | 3 | plasma plasminogen measurement | Acute ischaemic stroke: fibrinolytic treatment \| Thrombolysis in acute myocardial infarction \| Thrombolysis of occluded central venous access devices \| Thrombolytic treatment in acute massive pulmonary embolism | rs4252129 | 25208887 | 21442 | 1 | 1 | 1 |
| **DIO1** | PROPYLTHIOURACIL | Small molecule | 3 | thyroxine \|<br>thyroxine<br>measurement | Hyperthyroidism \| Thyrotoxic crisis \| Unlicensed product | rs2235544 | 23408906 | 1189 | 1 | 4 | 1 |
| **PDE4D** | DIPYRIDAMOLE | Small molecule | 4 | asthma | Alternative to exercise stress in thallium-201 myocardial imaging \| Ischemic stroke: Secondary prevention (with/without aspirin) \| Secondary prevention of ischaemic stroke \| Secondary prevention of transient ischaemic attacks \| Thromboembolism+prosthetic heart valve: prophylaxis (+oral anticoagulant) \| Transient ischemic attacks: Secondary prevention (with/without aspirin) | rs1588265 | 19426955 | 448153 | 1 | 2 | 1 |
| **ACHE** | RIVASTIGMINE | Small molecule | 4 | resting heart rate | Mild - moderate dementia in Alzheimer's disease \| Mild - moderate dementia in idiopathic Parkinson's disease | rs12666989 \|<br>rs314370 | 20639392 | 861 - 34407 | 3 - 7 | 9 | 4 |
| **ACHE** | NEOSTIGMINE<br>METHYLSULFATE | Small molecule | 4 | heart rate | Myasthenia gravis \| Paralytic ileus \| Paroxysmal supra-ventricular tachyarrhythmias \| Post operative distention \| Post operative urinary retention \| Reversal of residual competitive neuromuscular block \| Unlicensed product | rs13245899 | 23583979 | 861 - 34407 | 1 - 7 \| | 9 | 4 |
| **CHRM2** | TOLTERODINE<br>TARTRATE | Small molecule | 4 | heart rate | Symptomatic treatment of urinary urgency, frequency or urge incontinence | rs2350782 | 23583979 | 62368 | 1 | 3 | 1 |

Manual curation identified 1,523 discordant pairings of drug indications and disease associations, corresponding to 144 drug targets that were interpreted as plausible repurposing opportunities (Figure 4). After manual curation, uncertainty remained for 108 associations (52 targets) as to whether discordance represented a repurposing opportunity, or an unrecognised mechanism-based side effect. The remaining targets of licensed drugs mapped to LD intervals corresponding to GWAS traits unlikely to be of therapeutic interest (e.g. hair colour); or to a genetic association with a novel biomarker of uncertain biological function (e.g. a novel metabolite measured by a new metabolomics platform). Curators disagreed on the coding for GWAS associations corresponding to 4 licensed targets. For LD intervals corresponding to GWAS rediscoveries, the interval length was smaller, contained fewer genes, and the druggable gene was closer to the lead SNP than for those LD intervals where the indication and genetic association were discordant (Supplementary Table S2).

**Fig. 4.**
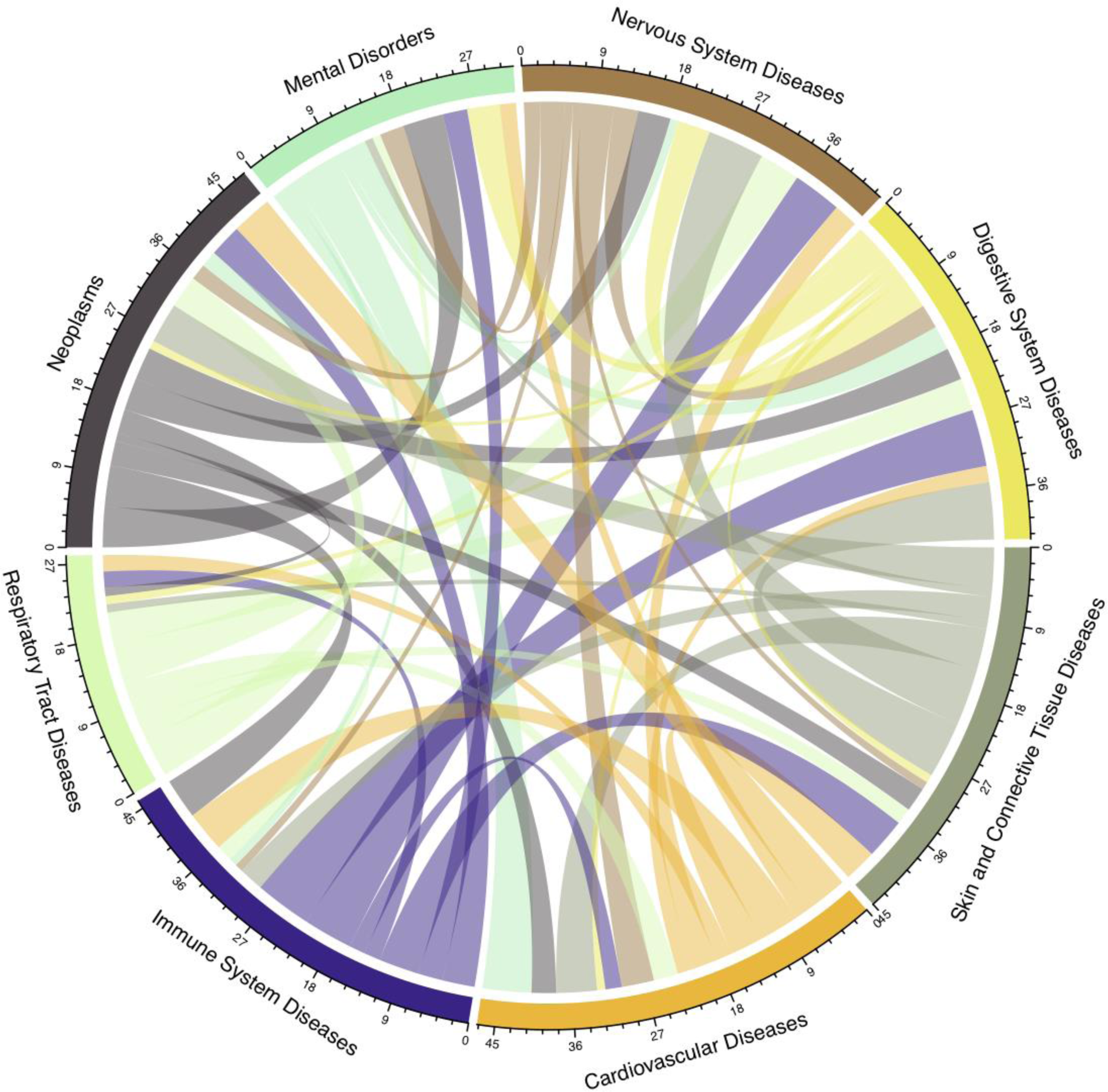
Potential repurposing opportunities from the discordant GWAS phenotype/drug indication matches (curation categories 5 and 6; see Methods). The disease categories on the circumference are MeSH root disease areas for drug indications and each unit on the scale represents a drug target gene. The chords connecting the disease areas represent drug targets for one disease area that are potential therapeutic targets in a different disease area. For this plot, only genes that overlapped a 50kbp flank surrounding the GWAS association are displayed to reduce the possibility discordance due to confounding by linkage disequilibrium.

### Translational opportunities unveiled by the data linkage

Figure 5 and Supplementary Figures S6 and S7 illustrate the result of mapping disease associations in the GWAS catalogue to the full set of druggable genes, the encoded proteins and allied compounds exhibiting binding affinity to these targets, regardless of development phase. For example, 84 studies in the GWAS catalogue reported findings pertaining to cardiovascular system diseases (39 disease sub categories), reporting 388 GWAS associations, mapping to 228 unique LD intervals containing 670 genes, of which 135 were in the druggable set. Of these, 29 genes were either the solitary occupant or one of only a pair of genes in the LD interval. We linked all 135 druggable genes identified in the cardiovascular category to 19,844 compounds with measured activities in ChEMBL (see Methods section Linking GWAS and drug target data), of which 512 had a United States Adopted Name (USAN) International Non-Proprietary Name (INN) or which were in late phase development, and 168 of which were previously licensed drugs. Based on comparisons between GWAS phenotype terms and treatment indications in the cardiovascular category, 8 drug target indications and genetic associations were concordant (target ‘rediscovery’) and 19 were discordant. Figure 6 illustrates the results of a similar mapping exercise for seven specific diseases (type 2 diabetes, hypertension, inflammatory bowel disease, asthma, coronary heart disease, schizophrenia, and Alzheimer’s disease).

**Fig. 5.**
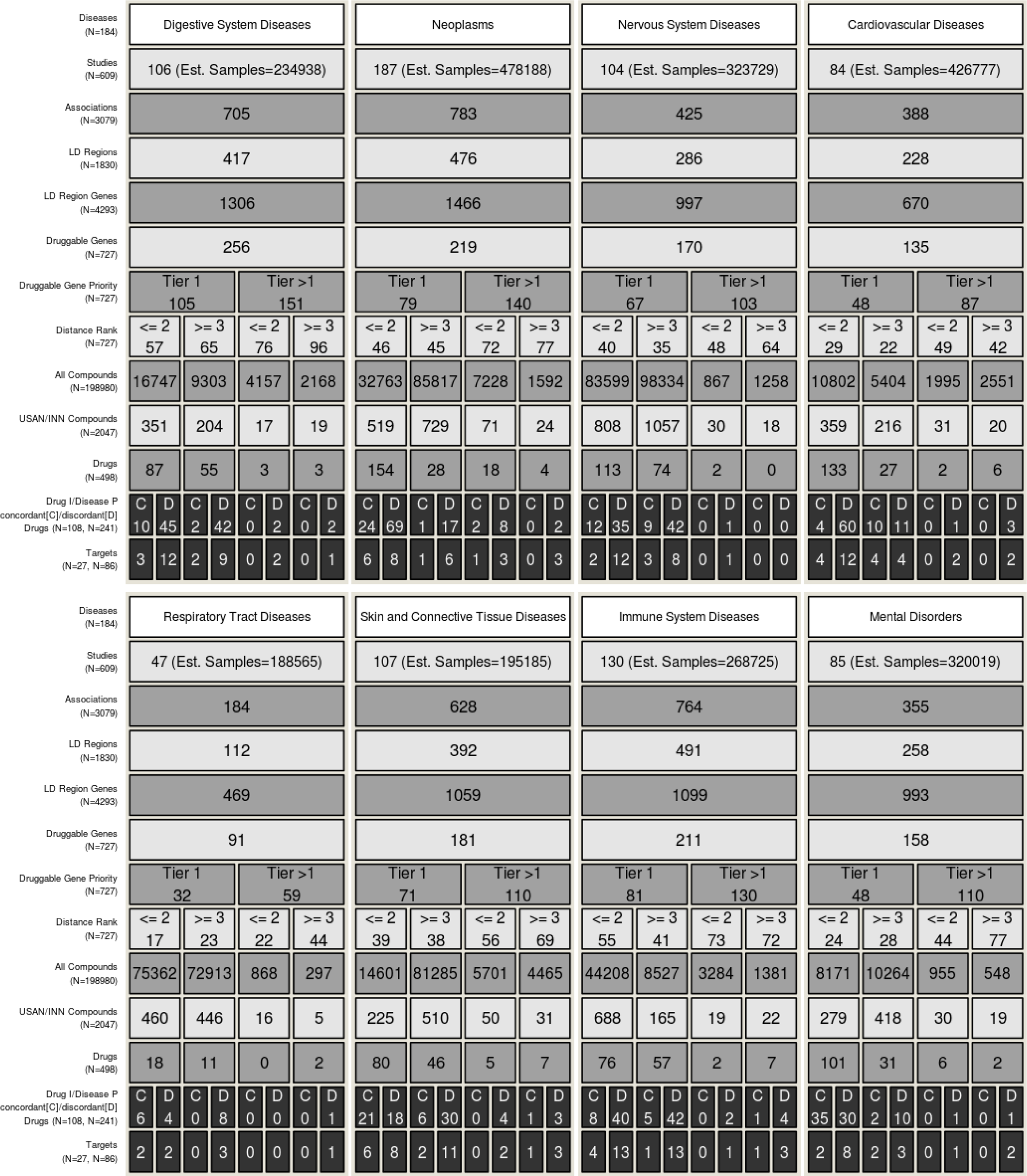
Translational potential for the top 8 most studied MeSH root disease areas (also see supplementary figure S1B. For each disease area, the figure illustrates the estimated number of GWAS, the number of associations (p≤5×10^-8^), the number of LD regions corresponding to those associations, the number of genes in those regions, and the number of those genes that encode druggable targets. Subsequent rows quantify the number of druggable genes by tier, and by distance rank from the GWAS SNP. The total number of compounds, compounds with an ISAN/INN and drugs corresponding to the druggable targets is also listed. In the penultimate row the numbers of drugs that that have an indication that is concordant (C) or discordant (D) with the GWAS phenotype are displayed. In the final row the number of cognate targets that for the concordant or discordant drugs are shown. Note that for the purposes of the figure a drug target is a single gene even if it is part of a complex that is targeted by the drug. Within each column the values are unique, i.e. number of unique associations (rsids). However, some values can be replicated across the figure, i.e. a GWAS study may have researched several of the disease areas. Additionally, there is some non-additivity between consecutive rows, namely Druggable Gene Priority - Distance Rank and Drugs - Drug indication/Disease Phenotypes. In the case of the former this is due to the same gene being further away from the associated variant in different studies, so falls into a different partition. For the later, this is due to missing indications for some of the drugs, so concordance or discordance could not be assigned. The values in the row labels represent the unique number of items across the row. The estimated number of samples is the sum of all the cases involved in the respective studies.

**Fig. 6.**
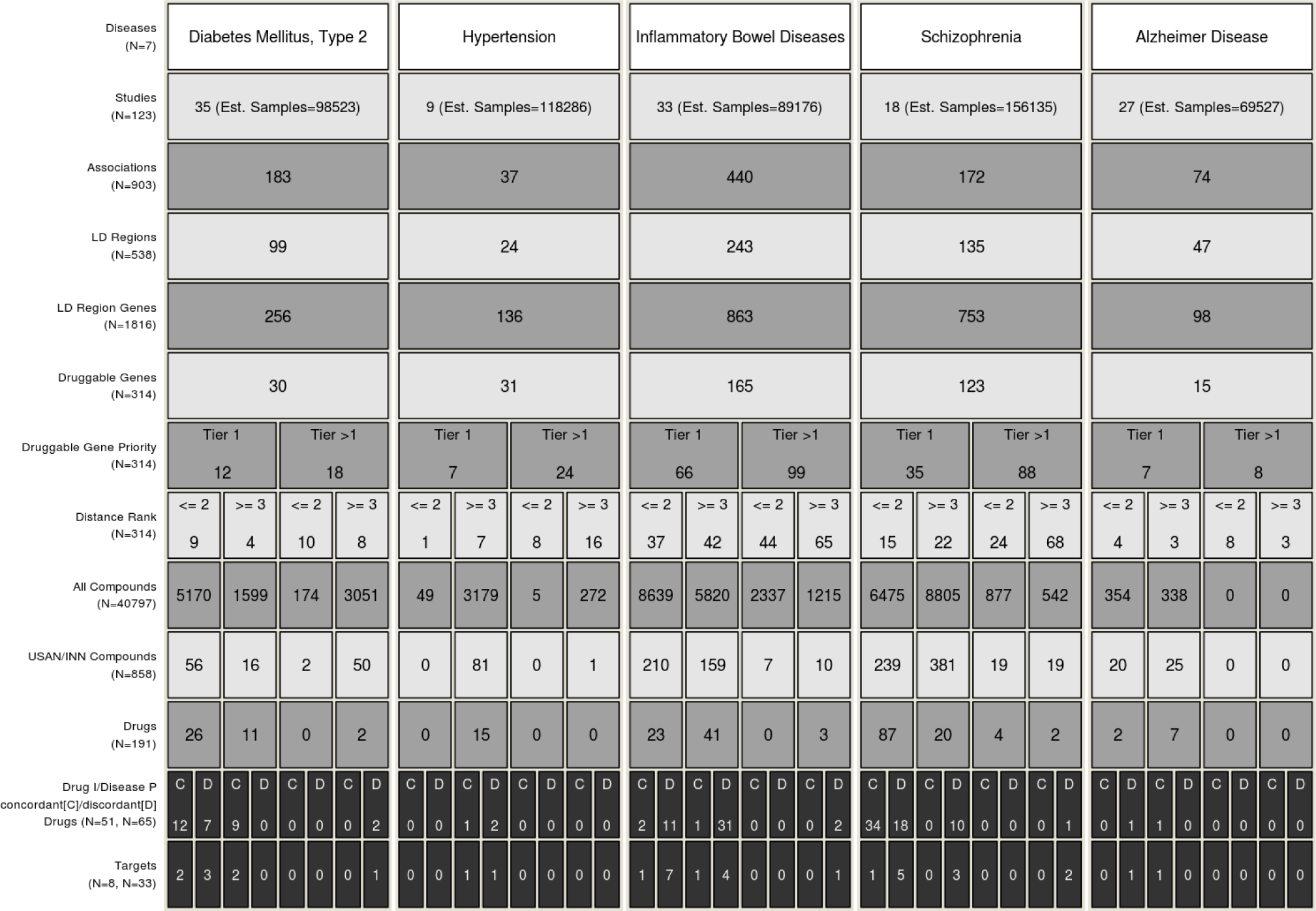
Translational potential for 5 specific diseases. Refer to Figure 5 legend for detailed explanation.

The proportion of druggable genes in LD intervals defined by GWAS SNPs for digestive system diseases (0.20, 95% CI: 0.12-0.27), neoplasms (0.15, 95%CI: 0.10-0.20), nervous system diseases (0.17, 95%CI: 0.10-0.24), cardiovascular diseases (0.20, 95%CI: 0.12-0.29), respiratory diseases (0.19, 95%CI: 0.08-0.31), skin and connective tissue diseases (0.17, 95%CI: 0.10-0.24), immune system diseases (0.19, 95%CI: 0.12-0.26) and mental health (0.16, 95%CI: 0.08-0.24) was similar to the proportion of druggable genes in the genome overall (4479/20,300 = 0.22).

### Coverage of the druggable genome by Illumina DrugDev and other widely used genotyping arrays

Capture of variation in druggable genes by the widely used genotyping arrays is illustrated in Figure 7, with reference to the 1000 genome European super population ancestry panels *(44)*. Disease-focused genotyping arrays and whole genome arrays with fewer than 600,000 SNPs used for many of the discoveries curated in the GWAS catalogue provided less comprehensive capture of variation in the druggable genome than the more recently developed arrays with several million SNPs (e.g. the Illumina Human Omni 2.5 Exome 8 and Illumina Omni 5). However, since no array to date has been designed specifically to ensure capture of variation in genes encoding druggable targets, we designed the content for an array (the Illumina DrugDev array) utilising the Illumina Infinium platform, that combines genome-wide tag SNP content of the Illumina Human Core array with 182,375 bespoke markers in 4479 druggable genes (see Methods). The median number of variants captured per kb of the druggable genome was very similar to that of the Illumina Human Omni 2.5 Exome 8 and Illumina Omni 5 (Figure 7 and Supplementary Figures S8 and S9) with an average of around 2.5 SNPs per kbp of the druggable genome, at an average of nearly 50 variants per gene array wide, with even denser coverage of Tier 1 and 2 genes.

**Fig. 7.**
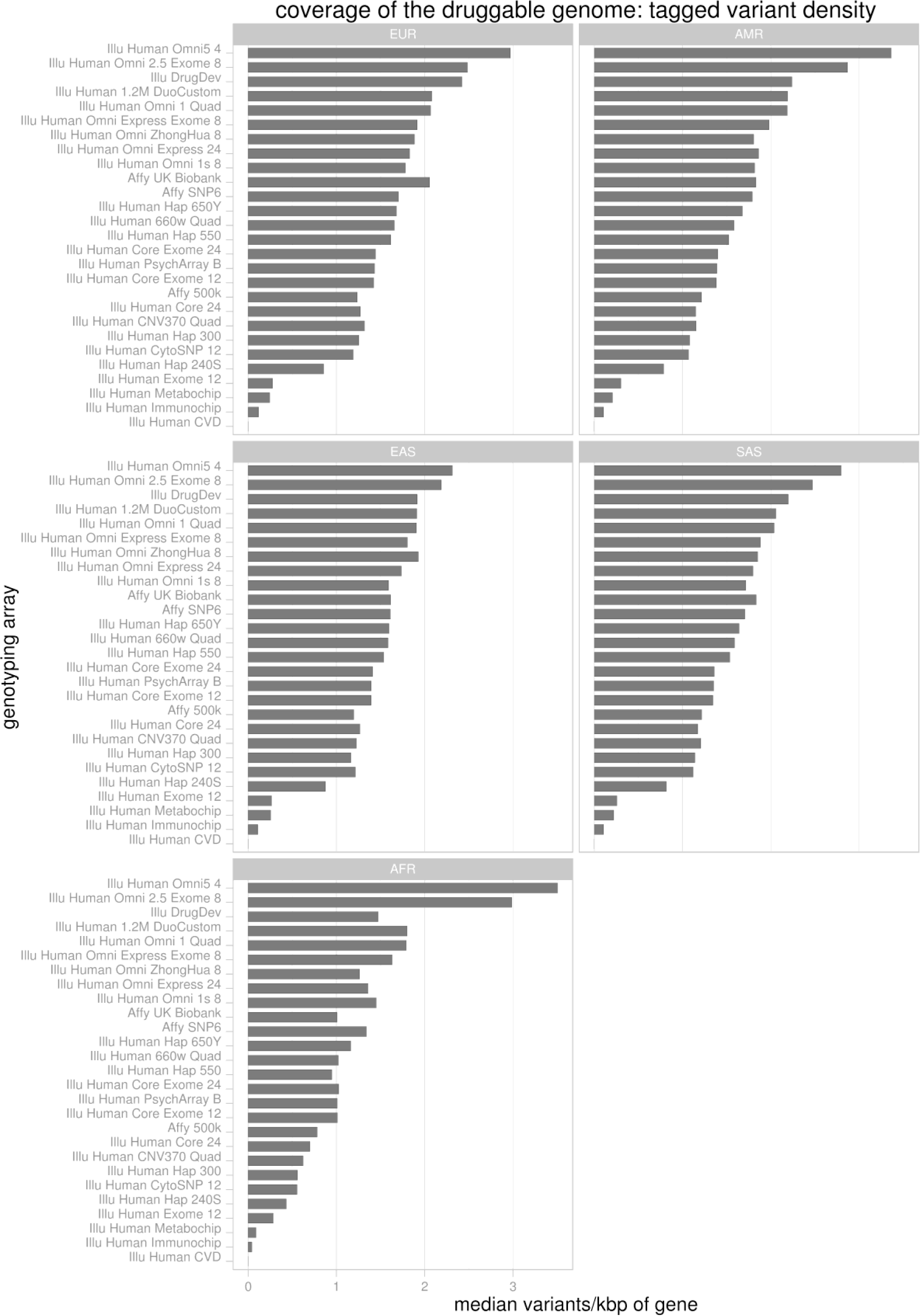
Tagged coverage of druggable genes in the 1000 genomes super populations. Coverage of the druggable gene set is represented as the median number of directly typed variants and variants in LD of *r^2^* ≥ 0.8 per kbp of druggable gene sequence. EUR – European, AMR - American, EAS – East Asian, SAS – South Asian, AFR – African.

All available genotyping arrays captured druggable genome variation most efficiently among European descent populations and most poorly among African descent populations (Figure 7 and Supplementary Figures S8 and S9). Outside of the European populations the high density Illumina Omni arrays gave superior coverage (for both directly genotyped variants and tagged variants) to all other genotyping arrays. The Affymetrix UK Biobank array displayed similar coverage to the Illumina DrugDev array in EUR populations but less complete coverage in non-European populations. A heat map summarising the coverage for each druggable gene, stratified by tier and 1000 genomes population groups, is shown in Figure 8. Results for tagged and directly typed variants in 1000 genomes sub-populations are shown in Supplementary Figure S10.

**Fig. 8.**
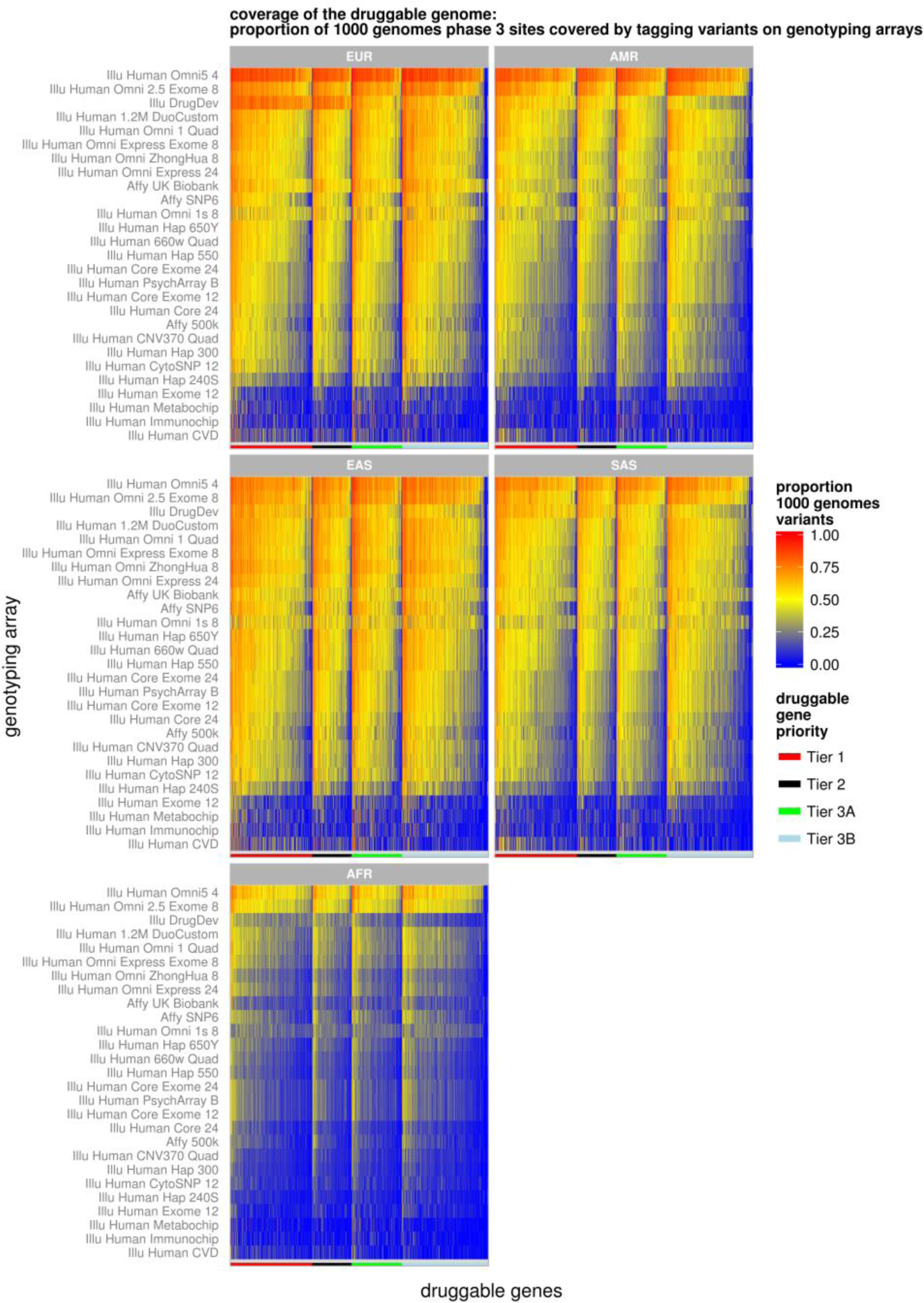
Tagged coverage of druggable genes in the 1000 genomes super populations. Coverage of the druggable gene set is represented as a proportion of 1000 genomes phase 3 variants (bialleilic with maf ≥ 0.005) that are either directly typed or in LD with *r^2^* ≥ 0.8 (tagged). Each column represents a druggable gene and each row a genotyping array. The druggable genes a grouped according to their druggability tier which is indicated by the colour bar at the base of each plot. To aid visualisation the druggable genes further are sorted within each tier on their median coverage across all the arrays and the genotyping arrays are sorted based on their median coverage of the druggable genome across all the 1000 genomes super populations. Note that all of the arrays contained some content that could not be mapped to the 1000 genomes phase 3 (see supplementary Figure S5). EUR – European, AMR - American, EAS – East Asian, SAS – South Asian, AFR – African.

## Discussion

By first re-estimating the boundaries of the druggable genome, and then mapping biomarker and disease associated loci from GWAS to genes encoding druggable targets, we demonstrate the extent to which GWAS have already rediscovered target-disease indications or mechanism-based adverse effects of licensed drugs. These findings indicate the potential of genetic association studies to systematically and accurately identify disease-specific drug targets across the spectrum of human diseases, addressing one of the key productivity limiting steps in drug development.

For example we found substantial potential for repurposing of drugs with licensed indications from one disease area to another (Figure 4), in keeping with previous analyses from the GWAS catalog that indicated that 17% of genes exhibit associations with more than one phenotype *(45)*. We also identified potential to progress or reposition compounds at earlier developmental stages, by mapping drug target loci implicated GWAS to the ChEMBL drug target annotations (Figure 5).

Despite the many novel therapeutic opportunities already arising from the mapping of existing genetic association findings to drug targets and compounds, there are strong reasons to suspect that the potential of this approach has yet to be maximised. Our analysis identified target-disease indication pairings (defined as a gene encoding a druggable target mapping to an LD interval containing a lead SNP from a GWAS) for 1,427 of the 4,479 druggable genes and 240 of the 652 genes encoding targets of licensed drugs. We might not have discovered associations for all genes in our druggable set because targets of drugs in development may truly play no role in any disease. However, alternative explanations are that only a fraction of diseases have been subjected to GWAS (451 out of 3022 conditions (the denominator is based on the number of bottom level MeSH disease areas)); that for many of the diseases that have been investigated by GWAS the sample sizes have been too small to detect all the responsible genes; or that there may have been incomplete coverage of certain druggable genes by the arrays most widely deployed in GWAS.

Genome wide association analyses continue to be published in new disease areas, and in new ethnic groups. Additional genetic discoveries are also being made with other types of array e.g. dense, locus-centric SNP arrays following up on GWAS findings that are currently not systematically captured by the GWAS catalog, eg. Cardiochip *(46)*, CardioMetabochip *(47)*, and Immunochip *(48)*, and by increases in sample size. Exome-arrays analyses are also unveiling rare, disease-associated variants under-represented in whole-genome arrays. Therefore, we anticipate that the current gap between druggable genes and GWAS findings will be reduced over time, particularly if such studies are extended to electronic health record datasets which form rich repositories of phenotypic traits and diagnostic codes.

Genetic profiling of a promising target against a range of outcomes can help evaluate the efficacy and safety of a target for the primary indication as well as the identification of additional disease indications to help plan drug development priorities. In order to stimulate the wider use of genetic association studies in drug development, and to ensure that such studies have comprehensive coverage of the druggable genome, we designed the content of a new array that combines focused coverage of the druggable genome within a whole genome scaffold. This array could be deployed to boost sample size and power in diseases already studied by GWAS to identify additional susceptibility loci and druggable targets. It could also help stimulate new druggable GWAS prioritised according to unmet therapeutic need. This would automatically lead to an abundance of target profiling information encompassing both efficacy and safety outcomes. This will need to be captured systematically, and curated consistently to help develop a repository of human drug targets linked to the predicted consequences of their pharmacological modification.

Some limitations of our analysis are noteworthy. The identification of repurposing opportunities in the current dataset relied on detecting discordance between a gene-disease association and the corresponding target-disease indication for a licensed drug, and excluding instances where this was likely to be due to a mechanism-based adverse effect. However, the lack of standardised vocabulary in licensing agency approval documents, and the scientific literature currently hampers this effort. We therefore used a combination of EFO and MeSH terms to harmonise nomenclature. Two qualified physicians then compared the annotations using a pre-specified classification system developed in a pilot study involving one fifth of the dataset. Greater efforts to harmonise terms both from the different ontologies (e.g. EFO, MeSH terms, the Disease Ontology (DO) and the Human Phenotype Ontology (HPO)) *(49–51)*, as well as from vocabularies for drug indications from the Anatomical Therapeutic Chemical (ATC) classification, electronic BNF and eMC+ terms would help generate standardised terminology to improve the efficiency of similar efforts in the future.

Where several genes occupy the same LD interval as a GWAS SNP, it may be difficult to determine which is causative. We took a pragmatic approach to this problem by classifying LD intervals containing druggable genes according to the total number of genes in the interval and the number and proximity of any druggable gene to the associated SNP. Approximately 529 unique LD intervals containing a variant with a significant association from a GWAS contained a single druggable gene. Such genes are strong positional candidates for the association. For the remainder, the LD interval included 2-146 genes (median 4 genes; excluding the 536 regions containing 0 genes, Figure 3), but a druggable gene was first or next most proximal gene to the association signal in 36.1% of these cases. The rediscovery of 183 target-indication or mechanism-based adverse pairings for licensed drugs using this indicates its validity of this approach. Previous Mendelian randomisation studies also provide reassurance that associations of SNPs in proximity to genes encoding druggable targets recapitulate the effects of drugs modifying the encoded protein pharmacologically *(13, 52, 18)*.

Nevertheless, we recognize that some misclassification is possible, for example when a causal signal arising from a gene encoding non-druggable protein occupies the same LD interval as a gene encoding a druggable target (confounding by linkage disequilibrium). Integrating information from feature annotation databases such as ENCODE *(53)* NIH Roadmap *(54)* and the Single Amino Acid Polymorphism Database (SAAP) *(55)* could help reduce misclassification. Localisation of causal genes could also be aided by evidence on the effect of genetic variants on the RNA transcription, on the activity or concentration of proteins and metabolites, combining new proteomic and metabolomics technologies that are scalable to large population studies *(56, 57)* with statistical approaches to assess whether association signals from the same region are consistent with the same causal variant *(58)*.

The Mendelian randomisation paradigm that underpins this strategy validates targets (within a defined disease context) and not compounds, although comparing the profile of effects of a genetic variant with those of a drug or developmental compound can help distinguish on- from off-target effects *(13, 18)*. For this reason RCTs will not be superseded by the approach we describe because any new molecule developed for a target of interest could have off-target actions that cannot be modelled genetically. Additionally, the effect of altering the level or function of a target may only be seen beyond some threshold, so that a weak genetic effect may not adequately model the effect of modifying the target pharmacologically *(26)*. Genetic evidence of a causal mechanism also does not guarantee its reversibility through pharmacological modification. For example, immune system related genetic variants associate with the risk of developing type I diabetes, but useful therapies arising from this knowledge may be difficult to realise because by the time the disease is diagnosed, immune mediated damage to the pancreatic beta-cells may be too advanced *(26)*. Despite these theoretical limitations, evidence is emerging that Mendelian randomisation studies have wide-ranging potential to improve the efficiency of drug development and reduce the risk of expensive late-stage failure.

In summary, we have shown an approach to focus and catalyse the use of genomic information to support drug target validation and which can be used to accurately match targets to disease indications and to identify rational repurposing opportunities for licensed drugs. The approach aligns well with proposals to ‘re-engineer’ translational science *(59)*. It could help address the efficiency and innovation problem and could serve as a basis for reinvigorating drug development through new academic-industry partnerships.

## Materials and Methods

### Assembly of a druggable gene set

The reference set of genes used to redefine the druggable genome comprised gene annotations from Ensembl v.73 with a biotype of ‘protein coding’. To this were added T-cell receptor and immunoglobulin genes, polymorphic pseudogenes, plus a number of additional genes that were annotated in Ensembl v.73 as non-protein coding but which were nevertheless believed to encode important proteins (e.g., SRD5A2, CYP4F8). Data were extracted via Biomart (http://www.ensembl.org/biomart). The content was assembled in three tiers:

*Tier 1* - This tier incorporated the targets of approved drugs and drugs in clinical development. Proteins that are targets of approved small molecule and biotherapeutics drugs were identified using manually curated efficacy target information from release 17 of the ChEMBL database *(60)*. An efficacy target was defined as the intended target for the drug as opposed to any other potential targets for which the drug shows high affinity binding. Where binding site information was available in ChEMBL, a non-drug-binding subunit of a protein complex were assigned to Tier 3, whereas the drug-binding subunit was included in Tier 1. Drugs in clinical development were identified from a number of sources: investor pipeline information from a number of large pharmaceutical companies (including Pfizer, Roche, GlaxoSmithKline, Novartis (oncology only), AstraZeneca, Sanofi, Lilly, Merck, Bayer and Johnson & Johnson – accessed June-August 2013) monoclonal antibody candidates and USAN applications from the ChEMBL database (release 17), and drugs in active clinical trials from clinicaltrials.gov *(61)*. Targets for these drug candidates were assigned from company pipeline information and scientific literature, where available. Where no reported target information could be found, a potential target was assigned through analysis of bioactivity data in ChEMBL, with the target having the highest dose-response measurement ≤ 100nM for the compound being assigned. All other human targets having an IC50/EC50/GI50/XC50/AC50/Kd/Ki/potency ≤100nM for an approved drug or USAN compound were also included in Tier1. Genes involved in ADME/drug disposition (phase I and II metabolic enzymes, transporters and modifiers) were identified from the PharmaADME.org extended set *(62)*.

*Tier 2* - This tier incorporated proteins closely related to drug targets or with associated drug-like compounds. Proteins closely related to targets of approved drugs were identified through a BLAST search (blastp) of Ensembl peptide sequences against the set of approved drug efficacy targets identified from ChEMBL previously *(38)*. Any genes where one or more Ensembl peptide sequences shared ≥50% identity (over ≥75% of the sequence) with an approved drug target were included. Putative targets with drug-like (Lipinski rule-of-five compliant) compounds having an IC50/EC50/GI50/XC50/AC50/Kd/Ki/potency ≤1µM were identified from ChEMBL and were also included in Tier 2.

*Tier 3* - This tier incorporated extracellular proteins and members of key drug-target families. Proteins distantly related to drug targets were identified through a BLAST search against the set of approved drug targets (as above), with any proteins sharing ≥25% identity over ≥75% of the sequence and with E-value ≤0.001 being included in the set. Members of five major ‘druggable’ protein families (GPCRs, kinases, ion channels, nuclear hormone receptors and phosphodiesterases) were extracted from KinaseSarfari *(63)*, GPCRSarfari *(64)* and IUPHARdb *(65)* and included in the Tier 3. Extracellular proteins were identified using annotation in UniProt *(66)* and Gene Ontology (GO) *(67)*. Since the potential size of the secreted/extracellular portion of the proteome (i.e., potential targets for monoclonal antibodies) is large, and the available number of markers for inclusion on the array was limited, this dataset was restricted to those proteins for which higher confidence annotations of extracellular localisation were available (not solely prediction of a signal peptide). Proteins annotated in UniProt as having a ‘secreted’ subcellular location, those containing a signal peptide, or those annotated as ‘Extracellular’ (where these annotations were supported by the following evidence types: experimental, probable, by_similarity) were included in Tier 3. Proteins annotated in GO with Cellular Component terms: GO:0005576 : extracellular region, GO:0005615 : extracellular space, GO:0005578 : proteinaceous extracellular matrix, GO:0031233 : intrinsic to external side of plasma membrane, GO:0031232 : extrinsic to external side of plasma membrane, GO:0071575 : integral to external side of plasma membrane, GO:0031362 : anchored to external side of plasma membrane, GO:0009897 : external side of plasma membrane, GO:0044214 : fully spanning plasma membrane, and supported by strong evidence (EXP, IDA, TAS), were also included in the tier. Finally, proteins known to be cluster of differentiation antigens (CD antigens), according to UniProt were also added to Tier 3. Since the final set of genes included in Tier 3 was large (2370 genes), this Tier was further subdivided to prioritise those genes that were in proximity (+/- 50Kb) to a GWAS SNP and had an extracellular location (Tier 3A). The remainder of the genes were assigned to Tier 3B.

### Pfam-A domain content

To evaluate the Pfam-A domain content for druggable genes, gene identifiers were converted to UniProt accession keys using the the UniProt web services *(66)*. Only UniProt accessions matching the regular expression pattern ‘[OPQ][0-9][A-Z0-9]{3}[0-9]’ were retained for further analysis. Pfam-A domains were extracted using the Xfam API *(68)*. For genes mapping to multiple UniProt accessions, we retained domain annotations for the UniProt accession mapping to the highest number of unique Pfam-A domains.

### Comparison of druggable gene sets

For comparison with genes covered on the Illumina DrugDev array, sets of druggable genes defined by Hopkins and Groom in 2002 and Russ and Lampel in 2005 were obtained from DGIdb. Gene names were converted to Ensembl gene identifiers using the Ensembl REST API *(69)*. The overlap between the three sets was determined and visualised using the Python module matplotlib_venn.

### Compilation of GWAS results

The GWAS catalog was downloaded from (http://www.ebi.ac.uk/gwas/api/search/downloads/alternative) on 21/07/2015. Several quality control and further post processing steps were then taken. The identifiers of associated variants were validated against Ensembl (version 79, build 37) using the perl API. This step returned the latest identifier and the build 37 coordinates; 707 associated variants could not be validated and were excluded. The GWAS catalog provides numerical effect estimates but does not specify the type of effect e.g odds ratio (OR) or beta co-efficient. Attempts were then made to resolve by utilising data in other fields (e.g. the presence of case or control in the discovery population fields) to classify the effect type as OR, beta or unknown. The discovery population field was also processed using a set of regular expressions to determine the sample size and populations used. The populations were then mapped to an appropriate 1000 genomes super population. Where no population name could be identified, EUR was used as a default as the majority of studies in the GWAS catalog were performed on Europeans. The pubmed identifier field was used to search pubmed using the Biopython API. MeSH terms for the publications were mapped to the association to provide structured phenotype descriptions. However, these study level descriptions may not apply to every association reported by the study, therefore the MeSH terms were manually curated for each association. These supplemented the experimental factor ontology terms (EFO) that are already present in the GWAS catalogue. Finally, the associations were filtered for those that are ≤ 5×10^-8^ so all data using in this study exceeded genome-wide significance.

### Assignment of LD intervals

The complete 1000 genomes phase 3 data (release 5) was downloaded from ftp://ftp.1000genomes.ebi.ac.uk/vol1/ftp/release/20130502. BCFTools (v1.2 using HTSlib 1.2.1) and used to subset the vcf files into sub- and super- population files *(70)*. For each population group, Plink v1.90b3d *(71)* was used to perform pairwise LD (*r^2^*) calculations between all variants in the processed GWAS catalog and bi-allelic 1000 genomes variants within a 1Mb flank either side of the GWAS variant having a maf ≥ 0.005. To reduce file size only *r^2^* values ≥ 0.2 were output. The extremities of the LD region surrounding each GWAS SNP were defined by the positions of the variants furthest upstream and downstream of this SNP with an *r^2^* value ≥ 0.5. Associated variants that were not present in the 1000 genomes panel that were not in LD with any other variants were given a nominal flank of 2.5Kb either size of the association.

### Linking GWAS and drug target data

Gene annotations were extracted from Ensembl version 79. After filtering out pseudogenes 38,352 genes remained. The set of genes was further reduced to those that overlapped an LD region surrounding an association. Within each associated LD region the absolute base pair distance of the closest point of a gene from the associated variant was calculated. Variants located within a gene were given a distance of 0bp. Genes were given a distance rank value according to their base pair distance. In the event of a distance rank tie, the gene with the oldest annotation date was assigned the lower rank.

Drug targets in ChEMBL 20 are annotated with UniProt accessions. The accessions were converted to Ensembl gene identifiers using the UniProt ID mapper (http://www.uniprot.org/uploadlists/). Drug target Ensembl gene IDs were then intersected with the IDs of genes within LD regions to give a set of drug targets in the proximity of associated variants.

### Evaluation of consistency between licensed drug indications and GWAS disease/biomarker traits

We evaluated the concordance between drug indication and disease association for those LD intervals defined by a GWAS SNP containing one or more genes encoding the target or targets of licensed drugs (Supplementary Figure S4). Two experienced clinicians used a pre-specified classification system developed in a pilot study of one-fifth of the total data set. Each physician was blinded to the identity of the gene encoding the druggable target within each LD interval. The outputs from the two physician-curators were then compared, any coding errors corrected, and inconsistencies between curators resolved by consensus, where agreement could be reached.

Category 0 referred to a situation where coding could not be completed because of missing data; 1 to a precise drug indication-target gene-disease association match; 2 to a drug indication-target gene-disease area association match; and 3 to a drug indication-target gene- mechanism-of-action association match. Categories 1 to 3 were defined as ‘concordant’. Category 4 referred to a drug mechanism based adverse effect-target gene-disease-association match; 5 to a drug indication-target gene-disease association mismatch with prior biological plausibility and 6 without prior biological plausibility; 7 to a trait unlikely to be of therapeutic interest (e.g. hair colour); and 8 to a genetic association with a novel biomarker of uncertain biological function (e.g. a metabolite measured by a metabolomics platform). For certain drug targets/genes, a 34 code was used to indicating that the genetic association finding could reflect both a mechanism of action and mechanism based adverse effect rediscovery. For example, the modification of certain electrocardiographic parameters by variants in the targets of certain antiarrhythmic drugs could reflect both their mechanism of action and the mechanism by such drugs produce their adverse effects. A 54 code was used when there was uncertainty about the direction of effect. A 9 code was assigned to the four cases where consensus could not be reached between the two curators. Categories 4, 5, 54, and 6 were referred to as discordant. Categories 1-4 and 34 were referred to collectively as ‘GWAS rediscoveries’ of known drug effects.

### Estimates of and confidence interval for the proportion of druggable genes in LD intervals

The proportion of druggable genes in LD intervals specified by GWAS associations in each MeSH disease or MeSH psychiatry category was calculated by dividing the number of druggable genes by the number of all genes with. 95% confidence intervals calculated assuming a binomial distribution, on the assumption that each study was independent.

### Design of the Illumina DrugDev Array and comparative analysis of coverage of variation in the druggable genome

*Selection of custom SNP content*

The design was based on three tiers, corresponding to the level of evidence for druggability of the encoded proteins, with highest priority given to genes in Tiers 1 and 2. Tag SNPs were selected from the 1000 genomes European ancestry populations (CEU/GBR/FIN/TSI). Associations (tagging) between SNPs were identified based on linkage disequilibrium (r2 >0.8). SNPs already covered, or tagged by the Human Core base content were not duplicated. Only SNPs with a minor allele frequency ≥1.5% were considered for inclusion. The tagging threshold was defined as the number of variants a SNP tags (including itself) and was varied according to the tier. For Tiers 1 and 2 a tagging threshold of 1 was applied, meaning that all SNPs were considered for inclusion, even if they only tag themselves. For Tier 3A a tagging threshold of 3, and for Tier 3B a threshold of 4 was used. SNPs were selected only if they were positioned within +/-2.5Kb of the druggable genes selected in the three tiers (defined as a region of 2.5Kb upstream of the Ensembl gene start position to 2.5Kb downstream of the Ensembl gene end position). SNPs from the Illumina Exome array were also included in the custom content where these were found within genes in Tiers 1, 2 and 3A. Again, any redundancy with the Human Core and selected tag SNP content was eliminated. A collection of mitochondrial tag SNPs from the Broad Institute, designed to capture common variation within the mitochondrial genome, were also included in the custom content ((http://www.broadinstitute.org/mpg/tagger/mito.html). This set comprises 64 SNPs, however only 56 of these loci were designable and included in the array. Finally, remaining space was filled with lead SNPs for any disease or trait association from the GWAS catalog, prioritising SNPs located within 50kb of a druggable gene, or within the gene boundaries of any protein-coding gene.

For Tier 1 genes, 99,102 custom markers were selected, including tag SNPs and HumanExome content. A further 17,944 of the HumanCore markers also fell within Tier 1 gene regions, giving 117,046 markers in total. Tier 2 included 40,943 custom markers and an additional 6,270 markers from the HumanCore fell within Tier 2 gene regions, resulting in a total of 47,213 markers. Genes in Tier 3 were represented by 38,858 custom markers. A further 21,626 HumanCore markers fell within Tier 3 gene regions, yielding 60,484 markers in total. In addition to coverage of genes encoding druggable targets, 6,400 SNPs associated with complex diseases or traits identified from the GWAS catalog and from selected gene-centric studies were also incorporated in the array content. Of these SNPs, 2,996 were already covered in the Human Core, or previously included in the custom content leaving 3,410 variants to be added (of which 1,395 were within Tier 1-3 gene regions). Finally, 53 mitochrondrial genome tag SNPs were also included, along with 9 mitochondrial genome exome SNPs. Considering all content, 226,138 markers were located in, or within +/-2.5 kb of, genes in the selected drugged, druggable and ADME sets. For the array as a whole, 78,175 markers were exonic, 286,577 intronic, and 27,393 located in 5’-, and 41,171 in 3’-untranslated regions respectively.

We used variants in the 1000 genomes phase 3 reference panel populations to compare coverage of the druggable genome by the new array and other commonly used genotyping arrays (see previous section). For this analysis, the variants on each array were first mapped to the 1000 genomes phase 3 reference panel and coverage then compared using two metrics: variant density (per kbp of the druggable gene) and the proportion of the variants in the druggable genome that were captured. We defined complete coverage of druggable genome as capture of all the bi-alleilic variants in a 1000 genomes phase 3 reference panel population with a minor allele frequency ≥ 0.005 (representing low frequency to common variants). Because of differences in variant content reported in successive genome builds, not all the content of the genotyping arrays could be mapped back to the 1000 genomes phase 3 reference set. However, the proportion of variants captured by each array that could be mapped to the 1000 genomes reference panel was very similar (Supplementary Figure S5).

### Evaluating genotyping array coverage of the DrugDev array

The build 37 genotyping array content for the Illumina arrays was downloaded from Will Rayner's array strand website (http://www.well.ox.ac.uk/∼wrayner/strand).Where multiple versions of an array exists the latest version number was downloaded. The Affymetrix array annotations were downloaded as SQLite databases from the Affymetrix website. 1000 genomes data was processed as described in the method for creating LD regions. Variants present on the genotyping arrays were mapped to 1000 genomes phase 3 using the following sequence: variants with rs identifiers were searched against the 1000 genomes sites file, if no match was obtained then synonyms of the rs identifier (obtained from Ensembl version 79 build 37) were searched. Variants not mapping by rs identifier were then mapped by chromosome, position and alleles (flipping the strand of the alleles where appropriate). Allele frequencies and variant tagging for each sub-population group were calculated using Plink(v1.90b3d *(72)*), tagging was restricted to bi-allielic low-frequency and common variants (maf ≥ 0.005) within 1Mb of the source SNP. Baseline 1000 genomes coverage of the druggable genome in the different sub-populations was ascertained using Bedtools (v2.22.1) to intersect 1000 genomes variants with a maf ≥ 0.005 against the druggable gene list (including 2.5 kbp up/down stream). Proportional coverage of the druggable genome by the different genotyping arrays was then ascertained by intersecting the baseline coverage with the 1000 genomes mapped array content.

### Indication and adverse effects of licensed therapies

Drug indication data was obtained from several sources. The primary source was the First Databank database (FDB, http://www.fdbhealth.co.uk/). This is a commercial database used by University College London Hospitals (UCLH) and a one off single release was kindly provided for research purposes by First Databank Europe Ltd. As FDB is used clinically this was regarded as the “gold standard” indication set used for the manual categorization of concordant/discordant drug/GWAS links (see above). FDB drug indications are tagged with Universal Medical Language System concept identifiers (CUIs) and could be mapped into MeSH and other ontologies within the UMLS meta-thesaurus *(49, 73)*. Drug indication data was obtained from ChEMBL 21. This was obtained by manual curation and mapping of data from FDA drug labels (https://dailymed.nlm.nih.gov/dailymed/), WHO ATC classification (http://www.whocc.no/atc_ddd_index/) and ClinicalTrials.gov (https://clinicaltrials.gov) This was used to supplement the FDB data and fill in indication data for drugs that were not present the FDB release.

Side effect data was obtained from the Side Effect Resource (SIDER) database *(74)*. The drug identifiers used in SIDER were mapped back to Chembl identifiers using a mapping file provided by SIDER. The side effects are provided as MedRA terms and UMLS CUIs and were mapped to MeSH terms using the UMLS.

## Acknowledgments

We thank Dr Cora Vacher and colleagues for helping to facilitate the design of the Illumina DrugDev Array. We would like to thank Dr. Reecha Sofat, Anita Jena-Smol for facilitating access to the First Databank Database and advice on designing database queries and First Databank Europe Ltd for providing a single copy of the database for research purposes.

## Author contributions

Chris Finan, Anna Gaulton, Felix Kruger, John Overington, Aroon Hingorani and Juan Pablo Casas developed the idea for the project and approaches to accurately connect genetic associations to drug targets and compounds. Anna Gaulton, Felix Kruger and John Overington updated estimates of the druggable genome. Luana Galver and Ryan Kelley worked with Anna Gaulton, Chris Finan, Tina Shah and Jorgen Engmann to develop SNP content for the Illumina DrugDev array. Anneli Karlsson curated target information for clinical stage drugs, Rita Santos curated target information for FDA approved drugs in the ChEMBL database, and Tom Lumbers and Aroon Hingorani compared indications and adverse effects of licensed drugs with disease associations from GWAS.

## Data and materials availability

Additional materials may be made available through contact with the authors.

## References

1. B. Munos, Lessons from 60 years of pharmaceutical innovation, Nat. Rev. Drug Discov. 8, 959–968 (2009).

2. S. M. Paul, D. S. Mytelka, C. T. Dunwiddie, C. C. Persinger, B. H. Munos, S. R. Lindborg, A. L. Schacht, How to improve R&D productivity: the pharmaceutical industry’s grand challenge, Nat. Rev. Drug Discov. 9, 203–214 (2010).

3. M. R. Macleod, A. Lawson McLean, A. Kyriakopoulou, S. Serghiou, A. de Wilde, N. Sherratt, T. Hirst, R. Hemblade, Z. Bahor, C. Nunes-Fonseca, A. Potluru, A. Thomson, J. Baginskitae, K. Egan, H. Vesterinen, G. L. Currie, L. Churilov, D. W. Howells, E. S. Sena, Risk of Bias in Reports of In Vivo Research: A Focus for Improvement, PLoS Biol 13, e1002273 (2015).

4. P. Perel, I. Roberts, E. Sena, P. Wheble, C. Briscoe, P. Sandercock, M. Macleod, L. E. Mignini, P. Jayaram, K. S. Khan, Comparison of treatment effects between animal experiments and clinical trials: systematic review., BMJ 334, 197 (2007).

5. H. B. van der Worp, D. W. Howells, E. S. Sena, M. J. Porritt, S. Rewell, V. O’Collins, M. R. Macleod, Can Animal Models of Disease Reliably Inform Human Studies?, PLoS Med 7, e1000245 (2010).

6. J. P. A. Ioannidis, Why Most Published Research Findings Are False, PLoS Med 2, e124 (2005).

7. D. Colquhoun, An investigation of the false discovery rate and the misinterpretation of p-values, R. Soc. Open Sci. 1, 140216 (2014).

8. L. G. Halsey, D. Curran-Everett, S. L. Vowler, G. B. Drummond, The fickle P value generates irreproducible results, Nat. Methods 12, 179–185 (2015).

9. H. Naci, J. P. A. Ioannidis, How Good Is “Evidence” from Clinical Studies of Drug Effects and Why Might Such Evidence Fail in the Prediction of the Clinical Utility of Drugs?, Annu. Rev. Pharmacol. Toxicol. 55, 169–189 (2015).

10. J. Arrowsmith, P. Miller, Trial Watch: Phase II and Phase III attrition rates 2011-2012, Nat. Rev. Drug Discov. 12, 569–569 (2013).

11. A. Hingorani, J. P. Casas, Interleukin-6 receptor pathways in coronary heart disease: a collaborative meta-analysis of 82 studies, The Lancet 379, 1205–1213 (2012).

12. A. Hingorani, S. Humphries, Nature’s randomised trials, The Lancet 366, 1906–1908 (2005).

13. R. Sofat, A. D. Hingorani, L. Smeeth, S. E. Humphries, P. J. Talmud, J. Cooper, T. Shah, M. S. Sandhu, S. L. Ricketts, S. M. Boekholdt, N. Wareham, K. T. Khaw, M. Kumari, M. Kivimaki, M. Marmot, F. W. Asselbergs, P. van Harst, R. P. F. Dullaart, G. Navis, D. J. van Veldhuisen, W. H. V. Gilst, J. F. Thompson, P. McCaskie, L. J. Palmer, M. Arca, F. Quagliarini, C. Gaudio, F. Cambien, V. Nicaud, O. Poirer, V. Gudnason, A. Isaacs, J. C. M. Witteman, C. M. van Duijn, M. Pencina, R. S. Vasan, R. B. D’Agostino, J. Ordovas, T. Y. Li, S. Kakko, H. Kauma, M. J. Savolainen, Y. A. Kesäniemi, A. Sandhofer, B. Paulweber, J. V. Sorli, A. Goto, S. Yokoyama, K. Okumura, B. D. Horne, C. Packard, D. Freeman, I. Ford, N. Sattar, V. McCormack, D. A. Lawlor, S. Ebrahim, G. D. Smith, J. J. P. Kastelein, J. Deanfield, J. P. Casas, Separating the Mechanism-Based and Off-Target Actions of Cholesteryl Ester Transfer Protein Inhibitors With CETP Gene Polymorphisms, Circulation 121, 52–62 (2010).

14. G. Davey Smith, Capitalizing on Mendelian randomization to assess the effects of treatments., J. R. Soc. Med. 100, 432–5 (2007).

15. J. P. Casas, E. Ninio, A. Panayiotou, J. Palmen, J. A. Cooper, S. L. Ricketts, R. Sofat, A. N. Nicolaides, J. P. Corsetti, F. G. R. Fowkes, I. Tzoulaki, M. Kumari, E. J. Brunner, M. Kivimaki, M. G. Marmot, M. M. Hoffmann, K. Winkler, W. März, S. Ye, H. A. Stirnadel, K.-T. Khaw, S. E. Humphries, M. S. Sandhu, A. D. Hingorani, P. J. Talmud, PLA2G7 Genotype, Lipoprotein-Associated Phospholipase A2 Activity, and Coronary Heart Disease Risk in 10 494 Cases and 15 624 Controls of European Ancestry, Circulation 121, 2284–2293 (2010).

16. CRP Coronary Heart Disease Genetics Collaboration, Association between C reactive protein and coronary heart disease: mendelian randomisation analysis based on individual participant data, BMJ 342, d548 (2011).

17. M. V. Holmes, T. Simon, H. J. Exeter, L. Folkersen, F. W. Asselbergs, M. Guardiola, J. a Cooper, J. Palmen, J. a Hubacek, K. F. Carruthers, B. D. Horne, K. D. Brunisholz, J. L. Mega, E. P. a van Iperen, M. Li, M. Leusink, S. Trompet, J. J. W. Verschuren, G. K. Hovingh, A. Dehghan, C. P. Nelson, S. Kotti, N. Danchin, M. Scholz, C. L. Haase, D. Rothenbacher, D. I. Swerdlow, K. B. Kuchenbaecker, E. Staines-Urias, A. Goel, F. van’t Hooft, K. Gertow, U. de Faire, A. G. Panayiotou, E. Tremoli, D. Baldassarre, F. Veglia, L. M. Holdt, F. Beutner, R. T. Gansevoort, G. J. Navis, I. Mateo Leach, L. P. Breitling, H. Brenner, J. Thiery, D. Dallmeier, A. Franco-Cereceda, J. M. a Boer, J. W. Stephens, M. H. Hofker, A. Tedgui, A. Hofman, A. G. Uitterlinden, V. Adamkova, J. Pitha, N. C. Onland-Moret, M. J. Cramer, H. M. Nathoe, W. Spiering, O. H. Klungel, M. Kumari, P. H. Whincup, D. a Morrow, P. S. Braund, A. S. Hall, A. G. Olsson, P. a Doevendans, M. D. Trip, M. D. Tobin, A. Hamsten, H. Watkins, W. Koenig, A. N. Nicolaides, D. Teupser, I. N. M. Day, J. F. Carlquist, T. R. Gaunt, I. Ford, N. Sattar, S. Tsimikas, G. G. Schwartz, D. a Lawlor, R. W. Morris, M. S. Sandhu, R. Poledne, A. H. Maitland-van der Zee, K.-T. Khaw, B. J. Keating, P. van der Harst, J. F. Price, S. R. Mehta, S. Yusuf, J. C. M. Witteman, O. H. Franco, J. W. Jukema, P. de Knijff, A. Tybjaerg-Hansen, D. J. Rader, M. Farrall, N. J. Samani, M. Kivimaki, K. a a Fox, S. E. Humphries, J. L. Anderson, S. M. Boekholdt, T. M. Palmer, P. Eriksson, G. Paré, A. D. Hingorani, M. S. Sabatine, Z. Mallat, J. P. Casas, P. J. Talmud, Secretory phospholipase A(2)-IIA and cardiovascular disease: a mendelian randomization study., J. Am. Coll. Cardiol. 62, 1966–76 (2013).

18. D. I. Swerdlow, D. Preiss, K. B. Kuchenbaecker, M. V. Holmes, J. E. L. Engmann, T. Shah, R. Sofat, S. Stender, P. C. D. Johnson, R. A. Scott, M. Leusink, N. Verweij, S. J. Sharp, Y. Guo, C. Giambartolomei, C. Chung, A. Peasey, A. Amuzu, K. Li, J. Palmen, P. Howard, J. A. Cooper, F. Drenos, Y. R. Li, G. Lowe, J. Gallacher, M. C. W. Stewart, I. Tzoulaki, S. G. Buxbaum, D. L. van der A, N. G. Forouhi, N. C. Onland-Moret, Y. T. van der Schouw, R. B. Schnabel, J. A. Hubacek, R. Kubinova, M. Baceviciene, A. Tamosiunas, A. Pajak, R. Topor-Madry, U. Stepaniak, S. Malyutina, D. Baldassarre, B. Sennblad, E. Tremoli, U. de Faire, F. Veglia, I. Ford, J. W. Jukema, R. G. J. Westendorp, G. J. de Borst, P. A. de Jong, A. Algra, W. Spiering, A. H. M. der Zee, O. H. Klungel, A. de Boer, P. A. Doevendans, C. B. Eaton, J. G. Robinson, D. Duggan, J. Kjekshus, J. R. Downs, A. M. Gotto, A. C. Keech, R. Marchioli, G. Tognoni, P. S. Sever, N. R. Poulter, D. D. Waters, T. R. Pedersen, P. Amarenco, H. Nakamura, J. J. V. McMurray, J. D. Lewsey, D. I. Chasman, P. M. Ridker, A. P. Maggioni, L. Tavazzi, K. K. Ray, S. R. K. Seshasai, J. E. Manson, J. F. Price, P. H. Whincup, R. W. Morris, D. A. Lawlor, G. D. Smith, Y. Ben-Shlomo, P. J. Schreiner, M. Fornage, D. S. Siscovick, M. Cushman, M. Kumari, N. J. Wareham, W. M. M. Verschuren, S. Redline, S. R. Patel, J. C. Whittaker, A. Hamsten, J. A. Delaney, C. Dale, T. R. Gaunt, A. Wong, D. Kuh, R. Hardy, S. Kathiresan, B. A. Castillo, P. van der Harst, E. J. Brunner, A. Tybjaerg-Hansen, M. G. Marmot, R. M. Krauss, M. Tsai, J. Coresh, R. C. Hoogeveen, B. M. Psaty, L. A. Lange, H. Hakonarson, F. Dudbridge, S. E. Humphries, P. J. Talmud, M. Kivimäki, N. J. Timpson, C. Langenberg, F. W. Asselbergs, M. Voevoda, M. Bobak, H. Pikhart, J. G. Wilson, A. P. Reiner, B. J. Keating, A. D. Hingorani, N. Sattar, HMG-coenzyme A reductase inhibition, type 2 diabetes, and bodyweight: evidence from genetic analysis and randomised trials, The Lancet 385, 351–361 (2014).

19. R. A. Scott, D. F. Freitag, L. Li, A. Y. Chu, P. Surendran, R. Young, N. Grarup, A. Stancáková, Y. Chen, T. V. Varga, others, A genomic approach to therapeutic target validation identifies a glucose-lowering GLP1R variant protective for coronary heart disease, Sci. Transl. Med. 8, 341ra76–341ra76 (2016).

20. P. Würtz, Q. Wang, P. Soininen, A. J. Kangas, G. Fatemifar, T. Tynkkynen, M. Tiainen, M. Perola, T. Tillin, A. D. Hughes, P. Mäntyselkä, M. Kähönen, T. Lehtimäki, N. Sattar, A. D. Hingorani, J.-P. Casas, V. Salomaa, M. Kivimäki, M.-R. Järvelin, G. Davey Smith, M. Vanhala, D. A. Lawlor, O. T. Raitakari, N. Chaturvedi, J. Kettunen, M. Ala-Korpela, Metabolomic Profiling of Statin Use and Genetic Inhibition of HMG-CoA Reductase, J. Am. Coll. Cardiol. 67, 1200–1210 (2016).

21. D. Melzer, J. R. B. Perry, D. Hernandez, A.-M. Corsi, K. Stevens, I. Rafferty, F. Lauretani, A. Murray, J. R. Gibbs, G. Paolisso, S. Rafiq, J. Simon-Sanchez, H. Lango, S. Scholz, M. N. Weedon, S. Arepalli, N. Rice, N. Washecka, A. Hurst, A. Britton, W. Henley, J. van de Leemput, R. Li, A. B. Newman, G. Tranah, T. Harris, V. Panicker, C. Dayan, A. Bennett, M. I. McCarthy, A. Ruokonen, M.-R. Jarvelin, J. Guralnik, S. Bandinelli, T. M. Frayling, A. Singleton, L. Ferrucci, A genome-wide association study identifies protein quantitative trait loci (pQTLs)., PLoS Genet. 4, e1000072 (2008).

22. R. Saxena, B. F. Voight, V. Lyssenko, N. P. Burtt, P. I. W. de Bakker, H. Chen, J. J. Roix, S. Kathiresan, J. N. Hirschhorn, M. J. Daly, T. E. Hughes, L. Groop, D. Altshuler, P. Almgren, J. C. Florez, J. Meyer, K. Ardlie, K. B. Boström, B. Isomaa, G. Lettre, U. Lindblad, H. N. Lyon, O. Melander, C. Newton-Cheh, P. Nilsson, M. Orho-Melander, L. Råstam, E. K. Speliotes, M.-R. Taskinen, T. Tuomi, C. Guiducci, A. Berglund, J. Carlson, L. Gianniny, R. Hackett, L. Hall, J. Holmkvist, E. Laurila, M. Sjögren, M. Sterner, A. Surti, M. Svensson, M. Svensson, R. Tewhey, B. Blumenstiel, M. Parkin, M. DeFelice, R. Barry, W. Brodeur, J. Camarata, N. Chia, M. Fava, J. Gibbons, B. Handsaker, C. Healy, K. Nguyen, C. Gates, C. Sougnez, D. Gage, M. Nizzari, S. B. Gabriel, G.-W. Chirn, Q. Ma, H. Parikh, D. Richardson, D. Ricke, S. Purcell, Genome-Wide Association Analysis Identifies Loci for Type 2 Diabetes and Triglyceride Levels, Science 316, 1331–1336 (2007).

23. L. J. Scott, K. L. Mohlke, L. L. Bonnycastle, C. J. Willer, Y. Li, W. L. Duren, M. R. Erdos, H. M. Stringham, P. S. Chines, A. U. Jackson, L. Prokunina-Olsson, C.-J. Ding, A. J. Swift, N. Narisu, T. Hu, R. Pruim, R. Xiao, X.-Y. Li, K. N. Conneely, N. L. Riebow, A. G. Sprau, M. Tong, P. P. White, K. N. Hetrick, M. W. Barnhart, C. W. Bark, J. L. Goldstein, L. Watkins, F. Xiang, J. Saramies, T. A. Buchanan, R. M. Watanabe, T. T. Valle, L. Kinnunen, G. R. Abecasis, E. W. Pugh, K. F. Doheny, R. N. Bergman, J. Tuomilehto, F. S. Collins, M. Boehnke, A Genome-Wide Association Study of Type 2 Diabetes in Finns Detects Multiple Susceptibility Variants, Science 316, 1341–1345 (2007).

24. D. Cook, D. Brown, R. Alexander, R. March, P. Morgan, G. Satterthwaite, M. N. Pangalos, Lessons learned from the fate of AstraZeneca’s drug pipeline: a five-dimensional framework, Nat. Rev. Drug Discov. 13, 419–431 (2014).

25. M. R. Nelson, H. Tipney, J. L. Painter, J. Shen, P. Nicoletti, Y. Shen, A. Floratos, P. C. Sham, M. J. Li, J. Wang, L. R. Cardon, J. C. Whittaker, P. Sanseau, The support of human genetic evidence for approved drug indications, Nat. Genet. advance online publication (2015), doi:10.1038/ng.3314.

26. R. M. Plenge, E. M. Scolnick, D. Altshuler, Validating therapeutic targets through human genetics, Nat. Rev. Drug Discov. 12, 581–594 (2013).

27. P. Sanseau, P. Agarwal, M. R. Barnes, T. Pastinen, J. B. Richards, L. R. Cardon, V. Mooser, Use of genome-wide association studies for drug repositioning, Nat. Biotechnol. 30, 317–320 (2012).

28. A. L. Hopkins, C. R. Groom, The druggable genome., Nat. Rev. Drug Discov. 1, 727–30 (2002).

29. A. P. Russ, S. Lampel, The druggable genome: an update., Drug Discov. Today 10, 1607–10 (2005).

30. R. D. Kumar, L.-W. Chang, M. J. Ellis, R. Bose, Prioritizing Potentially Druggable Mutations with dGene: An Annotation Tool for Cancer Genome Sequencing Data, PLoS ONE 8, e67980 (2013).

31. M. Griffith, O. L. Griffith, A. C. Coffman, J. V. Weible, J. F. McMichael, N. C. Spies, J. Koval, I. Das, M. B. Callaway, J. M. Eldred, C. A. Miller, J. Subramanian, R. Govindan, R. D. Kumar, R. Bose, L. Ding, J. R. Walker, D. E. Larson, D. J. Dooling, S. M. Smith, T. J. Ley, E. R. Mardis, R. K. Wilson, DGIdb: mining the druggable genome, Nat. Methods 10, 1209–1210 (2013).

32. M. Rask-Andersen, S. Masuram, H. B. Schiöth, The Druggable Genome: Evaluation of Drug Targets in Clinical Trials Suggests Major Shifts in Molecular Class and Indication, Annu. Rev. Pharmacol. Toxicol. 54, 9–26 (2014).

33. M. Whirl-Carrillo, E. M. McDonagh, J. M. Hebert, L. Gong, K. Sangkuhl, C. F. Thorn, R. B. Altman, T. E. Klein, Pharmacogenomics Knowledge for Personalized Medicine, Clin. Pharmacol. Ther. 92, 414–417 (2012).

34. F. Zhu, Z. Shi, C. Qin, L. Tao, X. Liu, F. Xu, L. Zhang, Y. Song, X. Liu, J. Zhang, B. Han, P. Zhang, Y. Chen, Therapeutic target database update 2012: a resource for facilitating target-oriented drug discovery, Nucleic Acids Res. 40, D1128–1136 (2012).

35. V. Law, C. Knox, Y. Djoumbou, T. Jewison, A. C. Guo, Y. Liu, A. Maciejewski, D. Arndt, M. Wilson, V. Neveu, A. Tang, G. Gabriel, C. Ly, S. Adamjee, Z. T. Dame, B. Han, Y. Zhou, D. S. Wishart, DrugBank 4.0: shedding new light on drug metabolism, Nucleic Acids Res. 42, D1091–1097 (2014).

36. A. Mullard, 2015 FDA drug approvals, Nat. Rev. Drug Discov. 15, 73–76 (2016).

37. L. a Hindorff, P. Sethupathy, H. a Junkins, E. M. Ramos, J. P. Mehta, F. S. Collins, T. a Manolio, Potential etiologic and functional implications of genome-wide association loci for human diseases and traits., Proc. Natl. Acad. Sci. U. S. A. 106, 9362–7 (2009).

38. A. P. Bento, A. Gaulton, A. Hersey, L. J. Bellis, J. Chambers, M. Davies, F. a Krüger, Y. Light, L. Mak, S. McGlinchey, M. Nowotka, G. Papadatos, R. Santos, J. P. Overington, The ChEMBL bioactivity database: an update., Nucleic Acids Res. 42, D1083–90 (2014).

39. First Databank Europe Ltd, Drug Data | FDB (First Databank) (available at http://www.fdbhealth.co.uk/).

40. A. J. Barr, Protein tyrosine phosphatases as drug targets: strategies and challenges of inhibitor development, Future Med. Chem. 2, 1563–1576 (2010).

41. S. Knapp, Emerging Target Families: Intractable Targets, Handb. Exp. Pharmacol. (2015), doi:10.1007/164_2015_28.

42. K. Eilbeck, S. E. Lewis, C. J. Mungall, M. Yandell, L. Stein, R. Durbin, M. Ashburner, The Sequence Ontology: a tool for the unification of genome annotations, Genome Biol. 6, R44 (2005).

43. M. Eijgelsheim, C. Newton-Cheh, N. Sotoodehnia, P. I. W. de Bakker, M. Müller, A. C. Morrison, A. V. Smith, A. Isaacs, S. Sanna, M. Dörr, P. Navarro, C. Fuchsberger, I. M. Nolte, E. J. C. de Geus, K. Estrada, S.-J. Hwang, J. C. Bis, I.-M. Rückert, A. Alonso, L. J. Launer, J. J. Hottenga, F. Rivadeneira, P. A. Noseworthy, K. M. Rice, S. Perz, D. E. Arking, T. D. Spector, J. A. Kors, Y. S. Aulchenko, K. V. Tarasov, G. Homuth, S. H. Wild, F. Marroni, C. Gieger, C. M. Licht, R. J. Prineas, A. Hofman, J. I. Rotter, A. A. Hicks, F. Ernst, S. S. Najjar, A. F. Wright, A. Peters, E. R. Fox, B. A. Oostra, H. K. Kroemer, D. Couper, H. Völzke, H. Campbell, T. Meitinger, M. Uda, J. C. M. Witteman, B. M. Psaty, H.-E. Wichmann, T. B. Harris, S. Kääb, D. S. Siscovick, Y. Jamshidi, A. G. Uitterlinden, A. R. Folsom, M. G. Larson, J. F. Wilson, B. W. Penninx, H. Snieder, P. P. Pramstaller, C. M. van Duijn, E. G. Lakatta, S. B. Felix, V. Gudnason, A. Pfeufer, S. R. Heckbert, B. H. C. Stricker, E. Boerwinkle, C. J. O’Donnell, Genome-wide association analysis identifies multiple loci related to resting heart rate, Hum. Mol. Genet. 19, 3885–3894 (2010).

44. The 1000 Genomes Project Consortium, A global reference for human genetic variation, Nature 526, 68–74 (2015).

45. S. Sivakumaran, F. Agakov, E. Theodoratou, J. G. Prendergast, L. Zgaga, T. Manolio, I. Rudan, P. McKeigue, J. F. Wilson, H. Campbell, Abundant Pleiotropy in Human Complex Diseases and Traits, Am. J. Hum. Genet. 89, 607–618 (2011).

46. B. J. Keating, S. Tischfield, S. S. Murray, T. Bhangale, T. S. Price, J. T. Glessner, L. Galver, J. C. Barrett, S. F. A. Grant, D. N. Farlow, H. R. Chandrupatla, M. Hansen, S. Ajmal, G. J. Papanicolaou, Y. Guo, M. Li, S. DerOhannessian, P. I. W. de Bakker, S. D. Bailey, A. Montpetit, A. C. Edmondson, K. Taylor, X. Gai, S. S. Wang, M. Fornage, T. Shaikh, L. Groop, M. Boehnke, A. S. Hall, A. T. Hattersley, E. Frackelton, N. Patterson, C. W. K. Chiang, C. E. Kim, R. R. Fabsitz, W. Ouwehand, A. L. Price, P. Munroe, M. Caulfield, T. Drake, E. Boerwinkle, D. Reich, A. S. Whitehead, T. P. Cappola, N. J. Samani, A. J. Lusis, E. Schadt, J. G. Wilson, W. Koenig, M. I. McCarthy, S. Kathiresan, S. B. Gabriel, H. Hakonarson, S. S. Anand, M. Reilly, J. C. Engert, D. A. Nickerson, D. J. Rader, J. N. Hirschhorn, G. A. FitzGerald, Concept, Design and Implementation of a Cardiovascular Gene-Centric 50 K SNP Array for Large-Scale Genomic Association Studies, PLoS ONE 3, e3583 (2008).

47. B. F. Voight, H. M. Kang, J. Ding, C. D. Palmer, C. Sidore, P. S. Chines, N. P. Burtt, C. Fuchsberger, Y. Li, J. Erdmann, T. M. Frayling, I. M. Heid, A. U. Jackson, T. Johnson, T. O. Kilpeläinen, C. M. Lindgren, A. P. Morris, I. Prokopenko, J. C. Randall, R. Saxena, N. Soranzo, E. K. Speliotes, T. M. Teslovich, E. Wheeler, J. Maguire, M. Parkin, S. Potter, N. W. Rayner, N. Robertson, K. Stirrups, W. Winckler, S. Sanna, A. Mulas, R. Nagaraja, F. Cucca, I. Barroso, P. Deloukas, R. J. F. Loos, S. Kathiresan, P. B. Munroe, C. Newton-Cheh, A. Pfeufer, N. J. Samani, H. Schunkert, J. N. Hirschhorn, D. Altshuler, M. I. McCarthy, G. R. Abecasis, M. Boehnke, The metabochip, a custom genotyping array for genetic studies of metabolic, cardiovascular, and anthropometric traits., PLoS Genet. 8, e1002793 (2012).

48. A. Cortes, M. A. Brown, Promise and pitfalls of the Immunochip, Arthritis Res. Ther. 13, 101 (2011).

49. F. B. Rogers, Communications to the Editor, Bull. Med. Libr. Assoc. 51, 114–116 (1963).

50. L. M. Schriml, C. Arze, S. Nadendla, Y.-W. W. Chang, M. Mazaitis, V. Felix, G. Feng, W. A. Kibbe, Disease Ontology: a backbone for disease semantic integration, Nucleic Acids Res. 40, D940–D946 (2012).

51. P. Robinson, S. Mundlos, The Human Phenotype Ontology, Clin. Genet. 77, 525–534 (2010).

52. The Interleukin-6 Mendelian Randomisation Analysis Consortium, The interleukin-6 receptor as a target for prevention of coronary heart disease: a mendelian randomisation analysis, The Lancet 379, 1214–1224 (2012).

53. I. Dunham, A. Kundaje, S. F. Aldred, P. J. Collins, C. a Davis, F. Doyle, C. B. Epstein, S. Frietze, J. Harrow, R. Kaul, J. Khatun, B. R. Lajoie, S. G. Landt, B.-K. Lee, F. Pauli, K. R. Rosenbloom, P. Sabo, A. Safi, A. Sanyal, N. Shoresh, J. M. Simon, L. Song, N. D. Trinklein, R. C. Altshuler, E. Birney, J. B. Brown, C. Cheng, S. Djebali, X. Dong, J. Ernst, T. S. Furey, M. Gerstein, B. Giardine, M. Greven, R. C. Hardison, R. S. Harris, J. Herrero, M. M. Hoffman, S. Iyer, M. Kelllis, P. Kheradpour, T. Lassman, Q. Li, X. Lin, G. K. Marinov, A. Merkel, A. Mortazavi, S. C. J. Parker, T. E. Reddy, J. Rozowsky, F. Schlesinger, R. E. Thurman, J. Wang, L. D. Ward, T. W. Whitfield, S. P. Wilder, W. Wu, H. S. Xi, K. Y. Yip, J. Zhuang, B. E. Bernstein, E. D. Green, C. Gunter, M. Snyder, M. J. Pazin, R. F. Lowdon, L. a L. Dillon, L. B. Adams, C. J. Kelly, J. Zhang, J. R. Wexler, P. J. Good, E. a Feingold, G. E. Crawford, J. Dekker, L. Elinitski, P. J. Farnham, M. C. Giddings, T. R. Gingeras, R. Guigó, T. J. Hubbard, M. Kellis, W. J. Kent, J. D. Lieb, E. H. Margulies, R. M. Myers, J. a Starnatoyannopoulos, S. a. Tennebaum, Z. Weng, K. P. White, B. Wold, Y. Yu, J. Wrobel, B. a Risk, H. P. Gunawardena, H. C. Kuiper, C. W. Maier, L. Xie, X. Chen, T. S. Mikkelsen, S. Gillespie, A. Goren, O. Ram, X. Zhang, L. Wang, R. Issner, M. J. Coyne, T. Durham, M. Ku, T. Truong, M. L. Eaton, A. Dobin, T. Lassmann, A. Tanzer, J. Lagarde, W. Lin, C. Xue, B. a Williams, C. Zaleski, M. Röder, F. Kokocinski, R. F. Abdelhamid, T. Alioto, I. Antoshechkin, M. T. Baer, P. Batut, I. Bell, K. Bell, S. Chakrabortty, J. Chrast, J. Curado, T. Derrien, J. Drenkow, E. Dumais, J. Dumais, R. Duttagupta, M. Fastuca, K. Fejes-Toth, P. Ferreira, S. Foissac, M. J. Fullwood, H. Gao, D. Gonzalez, A. Gordon, C. Howald, S. Jha, R. Johnson, P. Kapranov, B. King, C. Kingswood, G. Li, O. J. Luo, E. Park, J. B. Preall, K. Presaud, P. Ribeca, D. Robyr, X. Ruan, M. Sammeth, K. S. Sandu, L. Schaeffer, L.-H. See, A. Shahab, J. Skancke, A. M. Suzuki, H. Takahashi, H. Tilgner, D. Trout, N. Walters, H. Wang, Y. Hayashizaki, T. J. Hubbard, A. Reymond, S. E. Antonarakis, G. J. Hannon, Y. Ruan, P. Carninci, C. a Sloan, K. Learned, V. S. Malladi, M. C. Wong, G. P. Barber, M. S. Cline, T. R. Dreszer, S. G. Heitner, D. Karolchik, V. M. Kirkup, L. R. Meyer, J. C. Long, M. Maddren, B. J. Raney, L. L. Grasfeder, P. G. Giresi, B.-K. Lee, A. Battenhouse, N. C. Sheffield, K. a Showers, D. London, A. a Bhinge, C. Shestak, M. R. Schaner, S. K. Kim, Z. Z. Zhang, P. a Mieczkowski, J. O. Mieczkowska, Z. Liu, R. M. McDaniell, Y. Ni, N. U. Rashid, M. J. Kim, S. Adar, Z. Zhang, T. Wang, D. Winter, D. Keefe, V. R. Iyer, K. S. Sandhu, M. Zheng, P. Wang, J. Gertz, J. Vielmetter, E. C. Partridge, K. E. Varley, C. Gasper, A. Bansal, S. Pepke, P. Jain, H. Amrhein, K. M. Bowling, M. Anaya, M. K. Cross, M. a Muratet, K. M. Newberry, K. McCue, A. S. Nesmith, K. I. Fisher-Aylor, B. Pusey, G. DeSalvo, S. L. Parker, S. Balasubramanian, N. S. Davis, S. K. Meadows, T. Eggleston, J. S. Newberry, S. E. Levy, D. M. Absher, W. H. Wong, M. J. Blow, A. Visel, L. a Pennachio, L. Elnitski, H. M. Petrykowska, A. Abyzov, B. Aken, D. Barrell, G. Barson, A. Berry, A. Bignell, V. Boychenko, G. Bussotti, C. Davidson, G. Despacio-Reyes, M. Diekhans, I. Ezkurdia, A. Frankish, J. Gilbert, J. M. Gonzalez, E. Griffiths, R. Harte, D. a Hendrix, T. Hunt, I. Jungreis, M. Kay, E. Khurana, J. Leng, M. F. Lin, J. Loveland, Z. Lu, D. Manthravadi, M. Mariotti, J. Mudge, G. Mukherjee, C. Notredame, B. Pei, J. M. Rodriguez, G. Saunders, A. Sboner, S. Searle, C. Sisu, C. Snow, C. Steward, E. Tapanan, M. L. Tress, M. J. van Baren, S. Washieti, L. Wilming, A. Zadissa, Z. Zhengdong, M. Brent, D. Haussler, A. Valencia, A. Raymond, N. Addleman, R. P. Alexander, R. K. Auerbach, S. Balasubramanian, K. Bettinger, N. Bhardwaj, A. P. Boyle, A. R. Cao, P. Cayting, A. Charos, Y. Cheng, C. Eastman, G. Euskirchen, J. D. Fleming, F. Grubert, L. Habegger, M. Hariharan, A. Harmanci, S. Iyenger, V. X. Jin, K. J. Karczewski, M. Kasowski, P. Lacroute, H. Lam, N. Larnarre-Vincent, J. Lian, M. Lindahl-Allen, R. Min, B. Miotto, H. Monahan, Z. Moqtaderi, X. J. Mu, H. O’Geen, Z. Ouyang, D. Patacsil, D. Raha, L. Ramirez, B. Reed, M. Shi, T. Slifer, H. Witt, L. Wu, X. Xu, K.-K. Yan, X. Yang, Z. Zhang, K. Struhl, S. M. Weissman, S. a Tenebaum, L. O. Penalva, S. Karmakar, R. R. Bhanvadia, A. Choudhury, M. Domanus, L. Ma, J. Moran, A. Victorsen, T. Auer, L. Centarin, M. Eichenlaub, F. Gruhl, S. Heerman, B. Hoeckendorf, D. Inoue, T. Kellner, S. Kirchmaier, C. Mueller, R. Reinhardt, L. Schertel, S. Schneider, R. Sinn, B. Wittbrodt, J. Wittbrodt, G. Jain, G. Balasundaram, D. L. Bates, R. Byron, T. K. Canfield, M. J. Diegel, D. Dunn, A. K. Ebersol, T. Frum, K. Garg, E. Gist, R. S. Hansen, L. Boatman, E. Haugen, R. Humbert, A. K. Johnson, E. M. Johnson, T. M. Kutyavin, K. Lee, D. Lotakis, M. T. Maurano, S. J. Neph, F. V. Neri, E. D. Nguyen, H. Qu, A. P. Reynolds, V. Roach, E. Rynes, M. E. Sanchez, R. S. Sandstrom, A. O. Shafer, A. B. Stergachis, S. Thomas, B. Vernot, J. Vierstra, S. Vong, H. Wang, M. a Weaver, Y. Yan, M. Zhang, J. a Akey, M. Bender, M. O. Dorschner, M. Groudine, M. J. MacCoss, P. Navas, G. Stamatoyannopoulos, J. a Stamatoyannopoulos, K. Beal, A. Brazma, P. Flicek, N. Johnson, M. Lukk, N. M. Luscombe, D. Sobral, J. M. Vaquerizas, S. Batzoglou, A. Sidow, N. Hussami, S. Kyriazopoulou-Panagiotopoulou, M. W. Libbrecht, M. a Schaub, W. Miller, P. J. Bickel, B. Banfai, N. P. Boley, H. Huang, J. J. Li, W. S. Noble, J. a Bilmes, O. J. Buske, A. O. Sahu, P. V. Kharchenko, P. J. Park, D. Baker, J. Taylor, L. Lochovsky, An integrated encyclopedia of DNA elements in the human genome., Nature 489, 57–74 (2012).

54. B. E. Bernstein, J. A. Stamatoyannopoulos, J. F. Costello, B. Ren, A. Milosavljevic, A. Meissner, M. Kellis, M. A. Marra, A. L. Beaudet, J. R. Ecker, P. J. Farnham, M. Hirst, E. S. Lander, T. S. Mikkelsen, J. A. Thomson, The NIH Roadmap Epigenomics Mapping Consortium, Nat. Biotechnol. 28, 1045–1048 (2010).

55. N. S. Al-Numair, A. C. Martin, The SAAP pipeline and database: tools to analyze the impact and predict the pathogenicity of mutations, BMC Genomics 14, S4 (2013).

56. A. Lourdusamy, S. Newhouse, K. Lunnon, P. Proitsi, J. Powell, A. Hodges, S. K. Nelson, A. Stewart, S. Williams, I. Kloszewska, P. Mecocci, H. Soininen, M. Tsolaki, B. Vellas, S. Lovestone, R. Dobson, for the A. D. N. Initiative, Identification of cis-regulatory variation influencing protein abundance levels in human plasma, Hum. Mol. Genet. 21, 3719–3726 (2012).

57. K. Suhre, S.-Y. Shin, A.-K. Petersen, R. P. Mohney, D. Meredith, B. Wägele, E. Altmaier, P. Deloukas, J. Erdmann, E. Grundberg, C. J. Hammond, M. H. de Angelis, G. Kastenmüller, A. Köttgen, F. Kronenberg, M. Mangino, C. Meisinger, T. Meitinger, H.-W. Mewes, M. V. Milburn, C. Prehn, J. Raffler, J. S. Ried, W. Römisch-Margl, N. J. Samani, K. S. Small, H.-E. Wichmann, G. Zhai, T. Illig, T. D. Spector, J. Adamski, N. Soranzo, C. Gieger, Human metabolic individuality in biomedical and pharmaceutical research, Nature 477 (2011), doi:10.1038/nature10354.

58. C. Giambartolomei, D. Vukcevic, E. E. Schadt, L. Franke, A. D. Hingorani, C. Wallace, V. Plagnol, S. M. Williams, Ed. Bayesian Test for Colocalisation between Pairs of Genetic Association Studies Using Summary Statistics, PLoS Genet. 10, e1004383 (2014).

59. F. S. Collins, Reengineering Translational Science: The Time Is Right, Sci. Transl. Med. 3, 90cm17–90cm17 (2011).

60. A. Gaulton, L. J. Bellis, A. P. Bento, J. Chambers, M. Davies, A. Hersey, Y. Light, S. McGlinchey, D. Michalovich, B. Al-Lazikani, J. P. Overington, ChEMBL: a large-scale bioactivity database for drug discovery., Nucleic Acids Res. 40, D1100–7 (2012).

61. National Institute of Health, ClinicalTrials.gov ClinicalTrials.gov (available at https://clinicaltrials.gov/).

62. Montreal Heart Institute Pharmacogenomics Center, www.pharmaadme.org - Home (available at http://pharmaadme.org/joomla/index.php?option=com_frontpage&Itemid=1).

63. EMBL-EBI, Kinase SARfari (available at https://www.ebi.ac.uk/chembl/sarfari/kinasesarfari/).

64. EMBL-EBI, GPCR SARfari (available at https://www.ebi.ac.uk/chembl/sarfari/gpcrsarfari).

65. A. J. Pawson, J. L. Sharman, H. E. Benson, E. Faccenda, S. P. H. Alexander, O. P. Buneman, A. P. Davenport, J. C. McGrath, J. A. Peters, C. Southan, M. Spedding, W. Yu, A. J. Harmar, Nc-Iuphar, The IUPHAR/BPS Guide to PHARMACOLOGY: an expert-driven knowledgebase of drug targets and their ligands, Nucleic Acids Res. 42, D1098–D1106 (2014).

66. The Uniprot Consortium, UniProt: a hub for protein information, Nucleic Acids Res. 43, D204–D212 (2015).

67. M. Ashburner, C. A. Ball, J. A. Blake, D. Botstein, H. Butler, J. M. Cherry, A. P. Davis, K. Dolinski, S. S. Dwight, J. T. Eppig, M. A. Harris, D. P. Hill, L. Issel-Tarver, A. Kasarskis, S. Lewis, J. C. Matese, J. E. Richardson, M. Ringwald, G. M. Rubin, G. Sherlock, Gene ontology: tool for the unification of biology. The Gene Ontology Consortium., Nat. Genet. 25, 25–9 (2000).

68. R. D. Finn, A. Bateman, J. Clements, P. Coggill, R. Y. Eberhardt, S. R. Eddy, A. Heger, K. Hetherington, L. Holm, J. Mistry, E. L. L. Sonnhammer, J. Tate, M. Punta, Pfam: the protein families database, Nucleic Acids Res. 42, D222–D230 (2014).

69. A. Yates, K. Beal, S. Keenan, W. McLaren, M. Pignatelli, G. R. S. Ritchie, M. Ruffier, K. Taylor, A. Vullo, P. Flicek, The Ensembl REST API: Ensembl Data for Any Language, Bioinformatics, btu613 (2014).

70. P. Danecek, A. Auton, G. Abecasis, C. A. Albers, E. Banks, M. A. DePristo, R. E. Handsaker, G. Lunter, G. T. Marth, S. T. Sherry, G. McVean, R. Durbin, 1000 Genomes Project Analysis Group, The variant call format and VCFtools, Bioinformatics 27, 2156–2158 (2011).

71. S. Purcell, B. Neale, K. Todd-Brown, L. Thomas, M. a R. Ferreira, D. Bender, J. Maller, P. Sklar, P. I. W. de Bakker, M. J. Daly, P. C. Sham, PLINK: a tool set for whole-genome association and population-based linkage analyses., Am. J. Hum. Genet. 81, 559–75 (2007).

72. C. C. Chang, C. C. Chow, L. C. A. M. Tellier, S. Vattikuti, S. M. Purcell, J. J. Lee, Second-generation PLINK: rising to the challenge of larger and richer datasets, GigaScience 4 (2015), doi:10.1186/s13742-015-0047-8.

73. O. Bodenreider, The Unified Medical Language System (UMLS): integrating biomedical terminology, Nucleic Acids Res. 32, D267–D270 (2004).

74. M. Kuhn, I. Letunic, L. J. Jensen, P. Bork, The SIDER database of drugs and side effects, Nucleic Acids Res. 44, D1075–1079 (2016).

